# Tumor Cell Death Drives Tumor-Promoting IL-6⁺ iCAF formation via P2X7-activation

**DOI:** 10.64898/2026.03.18.712671

**Authors:** Ciarán McDonnell, Valeriya Zinina, Amina Othman, Larissa Launhardt, Anna Brichkina, Fatma Aktuna, Magdalena Brkic, Matthias Lauth, Eliana Stanganello, Mark Schmitt

## Abstract

Chemotherapy resistance in pancreatic ductal adenocarcinoma is commonly attributed to tumor cell–intrinsic mechanisms, yet how cytotoxic therapy reshapes the tumor microenvironment remains incompletely understood. Here we show that PDAC cells exposed to cytotoxic agents reprogram pancreatic stellate cells toward an inflammatory cancer-associated fibroblast phenotype. Mechanistically, chemotherapy triggers the release of ATP from dying PDAC cells, which activates P2X7 signaling in PSCs in a paracrine manner, leading ERK activation and inflammatory polarization. In turn, therapy-educated PSCs promote tumor cell proliferation, induce resistance-associated transcriptional programs and impair CD8⁺ T cell–mediated cytotoxicity in an IL-6–dependent manner. Pharmacological inhibition of P2X7 suppressed stromal IL-6 induction and enhanced gemcitabine efficacy in vivo. These findings identify a therapy-induced ATP–P2X7–IL-6 axis that links tumor cell death to stromal reprogramming and adaptive resistance in PDAC.

## Introduction

Pancreatic ductal adenocarcinoma (PDAC) is one of the most lethal malignancies worldwide with a 5-year relative survival rate of only around 11%, depending on tumor stage at the time of diagnosis. This poor prognosis is the result of the absence of symptoms in early stages, the consequent late diagnosis, as well as the limited effect of current therapies (1). PDAC is predominantly treated with chemotherapy, and despite initial responses, most tumors swiftly develop resistance leading to very poor outcomes of the current treatment regimens (2). While resistance mechanisms intrinsic to cancer cells have been extensively studied, increasing evidence indicates that therapy-induced alterations in the tumor microenvironment (TME) critically contribute to the failure of treatment (3).

A defining feature of the PDAC TME is its dense stroma, which is largely composed of cancer-associated fibroblasts (CAFs) derived from pancreatic stellate cells (PSCs)(4). CAFs are functionally heterogeneous and can adopt distinct phenotypic states, including myofibroblastic CAFs (myCAFs) and inflammatory CAFs (iCAFs)(5). In particular, iCAFs are characterized by the secretion of pro-inflammatory cytokines such as interleukin-6 (IL-6), which promote tumor cell proliferation, survival, immune evasion, and therapeutic resistance (6). Although CAF heterogeneity has been well described, the mechanisms by which cytotoxic therapy might shape fibroblast phenotypes within the PDAC microenvironment remains largely unexplored.

Cytotoxic chemotherapy induces tumor cell death accompanied by the release of danger-associated molecular patterns (DAMPs) and other soluble mediators. These factors can act in a paracrine manner to modulate neighboring viable cancer cells and stromal components (7). For example, we and others have previously demonstrated that therapy-induced DAMP release not only modulates anti-tumor immunity but also enhances intrinsic resistance of residual cancer cells (7, 8). However, whether dying PDAC cells similarly influence surrounding stromal fibroblasts and thereby contribute to a tumor-promoting microenvironment has not been elucidated.

In this study, we investigated how chemotherapy-treated PDAC cells affect PSC function and polarization. We show that factors released from dying tumor cells drive the polarization of PSCs toward an IL-6–secreting iCAF phenotype. Mechanistically, we identify extracellular ATP as a key mediator of this paracrine communication, acting through the purinergic receptor P2X7 to enhance IL-6 expression in PSCs. Functionally, PSCs polarized by dying tumor cells enhance PDAC cell proliferation, promote transcriptional programs associated with epithelial–mesenchymal transition and therapy resistance, and impair T cell–mediated tumor cell killing. Importantly, pharmacological inhibition of P2X7 improves the therapeutic efficacy of gemcitabine *in vivo*.

Together, our findings reveal a previously underappreciated consequence of chemotherapy in PDAC: the induction of a tumor-promoting, immunosuppressive stromal response driven by ATP/P2X7 signaling. Targeting this paracrine axis may represent a promising strategy to enhance cytotoxic therapy and counteract microenvironment-mediated resistance in PDAC.

## Results

### Chemotherapy-treated PDAC cells release factors polarizing pancreatic stellate cells to IL-6 secreting iCAFs

We have shown previously that cytotoxic therapy results in the release of danger-associated pattern molecules (DAMPs) from dying tumor cells that activate mTOR signaling in surrounding cancer cells to promote therapy resistance in colorectal and pancreatic cancer (8). Interestingly, when we analyzed PDAC tissues of gemcitabine treated KPC mice, we observed activation of mTOR signaling not only in cancer cells but also in surrounding α-SMA positive stromal cells (Fig. 1A), suggesting that fibroblasts might be similarly activated by tumor cell death. In order to examine whether pancreatic stromal cells are targets of factors released from dying tumor cells, we first treated murine KPC cells with 10 µM gemcitabine for 6 hours, which sufficed to initiate tumor cell death (Supplementary Data Fig. S1A-C). We then washed the cells with PBS to remove any excess gemcitabine and continued to culture the cells in media without gemcitabine for another 18 hours to allow the accumulation of factors released by dying tumor cells in the cell culture supernatants. We then used these supernatants (GEM-SU) or supernatants from vehicle treated cells (Control-SU) to compare their effects on pancreatic stellate cells (PSCs) -a main source of cancer-associated fibroblasts (CAFs) in PDAC (9) (Fig. 1B). Notably, the treatment with GEM-SU resulted in the activation of mTOR and MEK/ERK signaling in PSCs (Fig. 1C and D); which have both been described to enhance their tumor promoting phenotypes (10, 11). We next performed RNAseq analysis on PSCs treated with Control- and GEM-SU in order to compare how dying cancer cells alter the transcriptional profile of PSCs. Remarkably, culturing PSCs in supernatants of gemcitabine treated tumor cells increased their expression of iCAF related genes and downregulated the expression of myCAF markers, indicating that cytotoxic therapy indirectly promotes iCAF polarization via factors released from dying tumor cells (Fig. 1E). We next validated iCAF polarization via RT-qPCR analysis of known iCAF and myCAF related genes and observed that the GEM-SU treatment particularly increased the expression of secreted cytokines belonging to the Interleukin 6 (IL-6) family (i.e. IL-6, IL-11 and LIF) (Fig. 1F and G), which are well-known for their tumor-promoting effects in PDAC (12). Next, we confirmed the enhanced IL-6 expression in GEM-SU treated PSCs on the protein level via immunostaining (Fig. 1H and Supplementary Data Fig. S1D) and performed IL-6 specific ELISA-assays to demonstrate the increased release of IL-6 by PSCs exposed to GEM-SU (Fig. 1I). Of note, we detected similar activation of ERK-signaling and IL-6 production in human PSCs treated with supernatants of human PDAC cells exposed to cytotoxic agents (Fig. 1J and K and Supplementary Data Fig. S1E and F). As we detected increased activation of MEK/ERK and mTOR signaling in PSCs upon GEM-SU treatment, we next tested whether these pathways are required for the upregulation of IL-6 by GEM-SU. Notably, while the pharmacological inhibition of MEK prevented the upregulation of IL-6 in PSCs exposed to supernatants of gemcitabine treated KPCs, we did not observe any inhibitory effect using the mTOR antagonist Rapamycin (Fig. 1L and M and Supplementary Data Fig. S1G), indicating that IL-6 expression is regulated by MEK/ERK rather than mTOR signaling.

**Figure 1:**
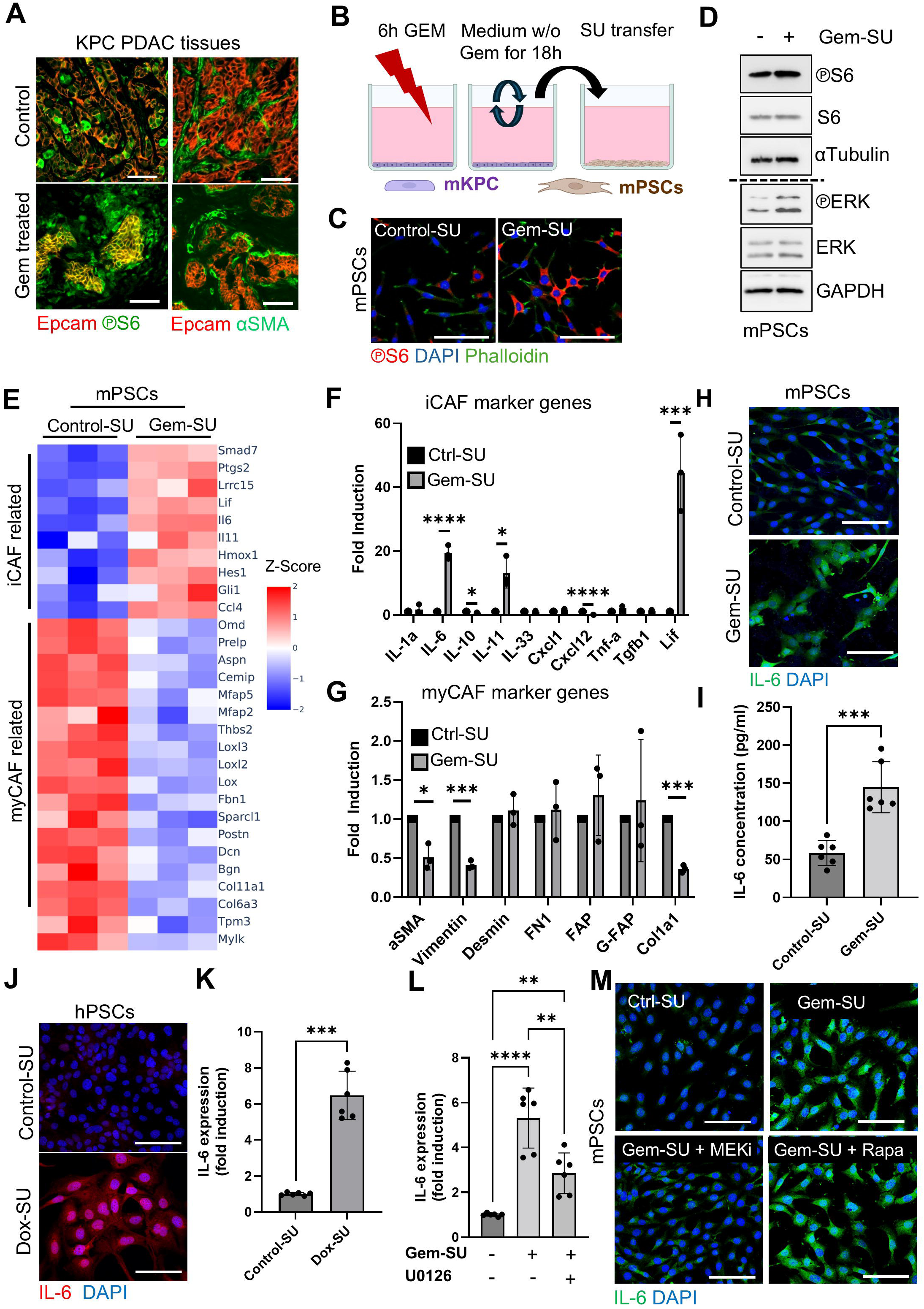
Gemcitabine treated PDAC cells polarize PSCs to IL-6 producing iCAFs. **A)** PDAC tissues of vehicle and gemcitabine treated KPC mice, stained for Epcam and phosphorylated S6 show activation of mTOR signaling in Epcam^+^ epithelial cells, as well as in adjacent Epcam^-^ stromal cells in gemcitabine treated mice (left panel). Epcam^-^ adjacent cells were positive for the fibroblast marker alpha-SMA (right panel). Representative images are shown (n =3 mice per group, scale bar = 100 µm). **B)** Scheme for the treatment of PSCs with supernatants of gemcitabine treated KPC cells (Gem-SU) supernatants of vehicle treated KPCs was used as control (Control-SU). **C)** Phospho-S6 staining of mPSCs treated with Control- or Gem-SU for 30 minutes. Cells were counterstained with Phalloidin, nuclei were counterstained with DAPI. Representative images are shown (n = 2 biological replicates, scale bar = 100 µm). **D)** Immunoblots of mPSC cells treated with Control- or Gem-SU for 30 min (n = 3 biological replicates). **E)** RNA-seq analysis of Control- or Gem-SU treated mPSCs. Heatmaps show differentially expressed iCAF- and myCAF related genes. Color codes indicate Z-scores of each condition (n=3 biological replicates). **F** and **G)** RT–qPCR analysis of the indicated genes in Control- and Gem-SU treated mPSCs (n = 3 (F) or 3 (G) biological replicates). **H)** Murine PSCs were treated with Control- and Gem-SU for 30 min and stained with an antibody against IL-6. Nuclei were counterstained with DAPI (scale bar = 100 µm, n= 3 biological replicates, quantification is shown in Supplementary Data Fig. S1D). **I)** IL-6 concentrations in Control- and Gem-SU, measured by ELISA (n = 3 biological replicates). **J)** Human PSCs were treated with supernatants of vehicle- (Control-SU) or doxorubicin treated MIA PaCa-2 cells (Dox-SU) for 30 min and stained with an antibody against IL-6. Nuclei were counterstained with DAPI (scale bar = 100 µm, n= 3 biological replicates, quantification is shown in Supplementary Data Fig. S1F). **K)** RT–qPCR analysis of IL-6 in human PSC cells treated for 24h with supernatants of vehicle (Control-SU) or doxorubicin treated (Dox-SU) MIA PaCa-2 cells (n = 3 biological replicates). **L)** IL-6 RT–qPCR analysis of mKPC treated with vehicle or gemcitabine in the presence or absence of the Mek-inhibitor U0126 (n = 3 biological replicates). **M)** Immunostaining against IL-6 in mPSCs treated with Control- or Gem-SU in the absence or presence of the Mek-inhibitor U0126 or the mTOR-inhibitor Rapamycin. Nuclei were counterstained with DAPI (scale bar = 100 µm, n= 3 biological replicates, quantification is shown in Supplementary Data Fig. S1G). All data are mean ± s.d. and were analyzed by two-tailed Student’s *t*-test (F,G,I,K), or one-way ANOVA (L) with Tukey’s multiple comparison test (*p≤0.05; **p≤0.01; ***p≤0.0005; **** p≤0.0001; only p values ≤ 0.05 are shown). Illustration in (B) was created using BioRender.com.

### Dying PDAC cells release ATP to stimulate IL-6 expression of PSCs via activation of P2X7

Our next aim was to reveal which paracrine factors released by dying tumor cells augment IL-6 production in PSCs. We have shown previously that in response to cytotoxic therapy dying tumor cells release ATP, to enhance therapy resistance by activating purinergic signaling in surrounding cancer cells (8) and interestingly, it has been described that purinergic signaling can increase IL-6 expression in PSCs (13). In line with a potential involvement of purinergic signaling, we observed an upregulation of genes regulated by purinergic signaling in PSCs exposed to GEM-SU, as well as an increase of extracellular ATP in supernatants of gemcitabine treated KPCs (Fig. 2A and Supplementary Data Fig. S2A). Hence, we examined whether ATP contributes to the increased IL-6 expression in GEM-SU treated PSCs. To do so, we first tested whether ATP is capable of activating MAPK-signaling and IL-6 expression in PSCs and detected increased phosphorylation of ERK and upregulated expression of IL-6 in PSCs treated with ATP, resembling the effect of supernatants of gemcitabine treated KPCs (Fig. 2B and C). In order to confirm that ATP released from gemcitabine treated PSCs is required for the paracrine activation of PSCs, we pre-treated GEM-SU with Apyrase to hydrolyze ATP, which prevented GEM-SU to induce ERK-signaling in PSCs (Fig. 2D).

**Figure 2:**
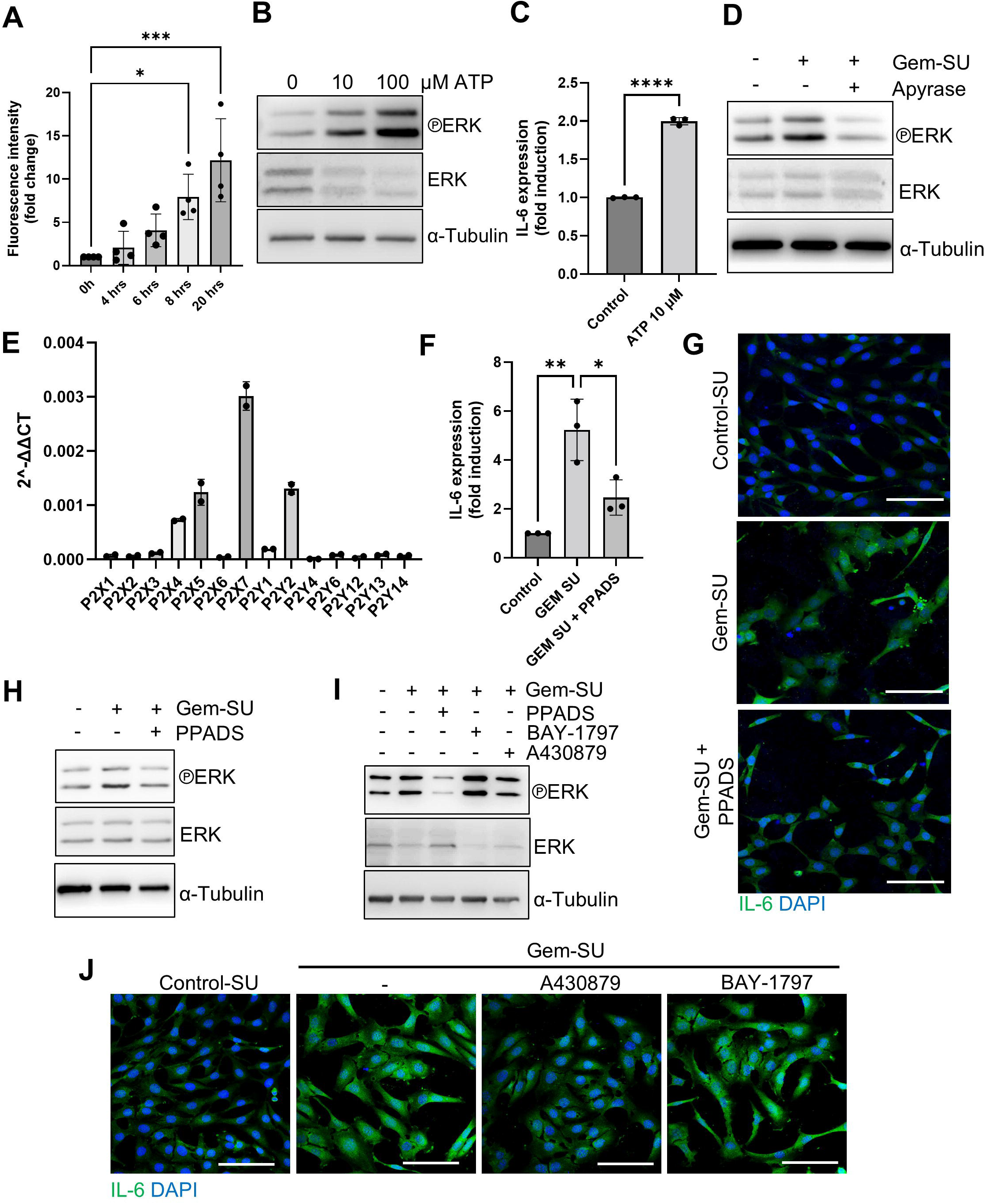
Dying PDAC induce IL-6 expression in PSCs via activation of ATP/P2X7-signaling. **A)** ATP level in supernatants of gemcitabine treated mKPCs (n = 3 biological replicates). **B)** Immunoblots of mPSC cells treated with indicated concentrations of ATP for 15 min (n = 3 biological replicates). **C)** IL-6 RTqPCR analysis of mPSCS treated with 10 µM ATP for 24 hrs. **D)** Immunoblots of mPSC cells treated with Control-SU, Gem-SU or Gem-SU pre-treated with Apyrase (n = 3 biological replicates). **E)** RT–qPCR analysis of the indicated genes in mPSCs (n= 2 biological replicates). **F)** RT-qPCR analysis of IL-6 expression in mPSCs treated for 24h with Control-SU or Gem-SU in the absence or presence of PPADS. **H)** IL-6 immunostaining of mPSCs treated with Control- or Gem-SU in the absence or presence of P2R-inihibitor PPADS. Nuclei were counterstained with DAPI (scale bar = 100 µm, n= 3 biological replicates, quantification is shown in Supplementary Data Fig. S2B). **H)** Immunoblots of mPSC cells treated with Control- or Gem-SU in the absence or presence of the P2-inhibitor PPADS (n = 3 biological replicates). **I)** Immunoblots of mPSC cells treated with Control- or Gem-SU in the absence or presence of the inhibitors against P2X (PPADS), P2X7 (A430879), P2X4 (BAY-1797) (n = 3 biological replicates). **J)** IL-6 immunostaining of mPSCs treated with Control- or Gem-SU in the absence or presence of inhibitors against P2X7 (A430879) and P2X4 (BAY-1797) (scale bar = 100 µm, n= 3 biological replicates, quantification is shown in Supplementary Data Fig. S2C). All data are mean ± s.d. and were analyzed by two-tailed Student’s *t*-test (C) or one-way ANOVA (A, F) with Tukey’s multiple comparison test (*p≤0.05; **p≤0.01; ***p≤0.0005; **** p≤0.0001; only p values ≤ 0.05 are shown).

The cell surface purinoreceptors responding to ATP encompass seven P2X ligand-gated ion-channels and seven G-protein coupled P2Y receptors (14) and we performed qRT-PCRs to analyze their expression pattern in mPSCs. According to our results, mPSCs mainly express the P2X receptors P2X4, P2X5 and P2X7 and the P2Y receptor P2Y2 (Fig. 2E). Particularly P2X receptors have been shown to promote IL-6-expression in PSCs (13) and hence we tested whether polarization of PSCs to IL-6 expressing iCAFs is dependent on P2X receptor activation by supplementing the supernatants of gemcitabine treated KPCs with the non-selective P2X receptor antagonist pyridoxalphosphate-6-azophenyl-2’,4’-disulfonic acid (PPADS) (15), which markedly inhibited the upregulation of IL-6 by GEM-SU on the RNA and protein level (Fig. 2F-H and Supplementary Data Fig. S2B). Next, we addressed which specific P2X-receptors are involved in polarization of PSCs. For this, we treated PSCs with GEM-SU either in the absence or presence of the non-selective P2R inhibitor PPADS, or the selective P2X4 Inhibitor BAY-1797 or the selective P2X7 inhibitor A438079, respectively (16). Due to the lack of a specific P2X5 inhibitor we could not address the specific blockade of P2X5. As in the previous experiment, the blockade of all P2X receptors by PPADS strongly inhibited ERK activation by GEM-SU. Regarding the selective P2X antagonists, the inhibition of P2X7 showed a higher capacity to counteract ERK activation when compared to the inhibition of P2X4 (Fig. 2I). Likewise, the inhibition of P2X7 was more potent in inhibiting IL-6 upregulation in PSCs exposed to GEM-SU when compared to P2X4 blockade (Fig. 2J and Supplementary Data Fig. S2C), corroborating a key role of P2X7 in the tumor cell death-induced polarization of PSCs. Moreover, PPADS and A438079 blocked ERK activation and decreased IL-6 production of human PSCs exposed to supernatants from doxorubicin treated MIA-PaCa-2 cells (Supplementary Data Fig. S2D and E). Notably, the inhibition of P2X7 not only suppressed the capacity of GEM-SU to promote the expression of IL-6 but also mitigated the expression of other IL-6 family members in GEM-SU treated PSCs (Supplementary Data Fig. S2F).

### PSCs polarized by dying tumor cells promote cancer cell proliferation via IL-6

Inflammatory CAFs are well known for their protumorigenic effects on PDAC cells (18). We therefore tested whether PSCs, polarized by GEM-SU, in turn release factors promoting tumor cell growth. To this aim, we first exposed PSCs as described previously to Control- or GEM supernatants for 6 hours. We then washed and cultured the PSCs in fresh culture medium for another 18 hours in order to enrich the supernatants for factors released by polarized PSCs and collected these supernatants (GEM-PSC-SU) for the subsequent treatment of KPCs (Fig. 3A). As control we used supernatants from PSCs exposed to culture medium of vehicle treated KPCs (Control-PSC-SU). Strikingly, KPCs exposed to GEM-PSC-SU, showed an increased expression of the proliferation marker Ki67 when compared to KPCs exposed to Control-PSC-SU, which was significantly reduced when either PSC cells where treated with A430879 when exposed to GEM-SU or when KPCs were treated with the IL-6-receptor antagonist LMT-28 during the treatment with GEM-PSC-SU (Fig. 3B), demonstrating that the growth promoting effects of PSCs is on the one hand dependent on their polarization via P2X7 and on the other hand on their release of IL-6. In line with elevated Ki67 mRNA, we observed an increase of PDAC cells with nuclear staining of Ki67 in both murine and human GEM-PSC-SU treated cells, which was blocked when PDAC cells were treated with the IL-6- receptor antagonist LMT-28(19) (Fig. 3C and D). Of note, the increase of nuclear Ki67 was also reduced by treating PSCs with the P2X7 inhibitor A438079 (Supplementary Data Fig. S3A). Besides the increased expression of the proliferation marker Ki67, we observed enhanced PDAC tumoroid growth upon exposure to GEM-PSC-SU when compared to tumoroids treated with control PSC supernatants (Fig. 3E). Also, here we could counteract this increase by inhibiting P2X7 signaling in PSCs or by inhibiting the activation of IL-6-receptors in KPCs (Fig. 3E). We furthermore performed bulk RNAseq analysis on KPC cells cultured in Control- or Gem-PSC-SU, which not only confirmed an increase of proliferation marker genes upon Gem-PSC-SU treatment but also increased expression of EMT-related genes and genes involved in therapeutic resistance (Supplementary Data Fig. S3B and C), indicating that the tumor promoting function of polarized PSCs goes beyond enhancing tumor cell proliferation.

**Figure 3:**
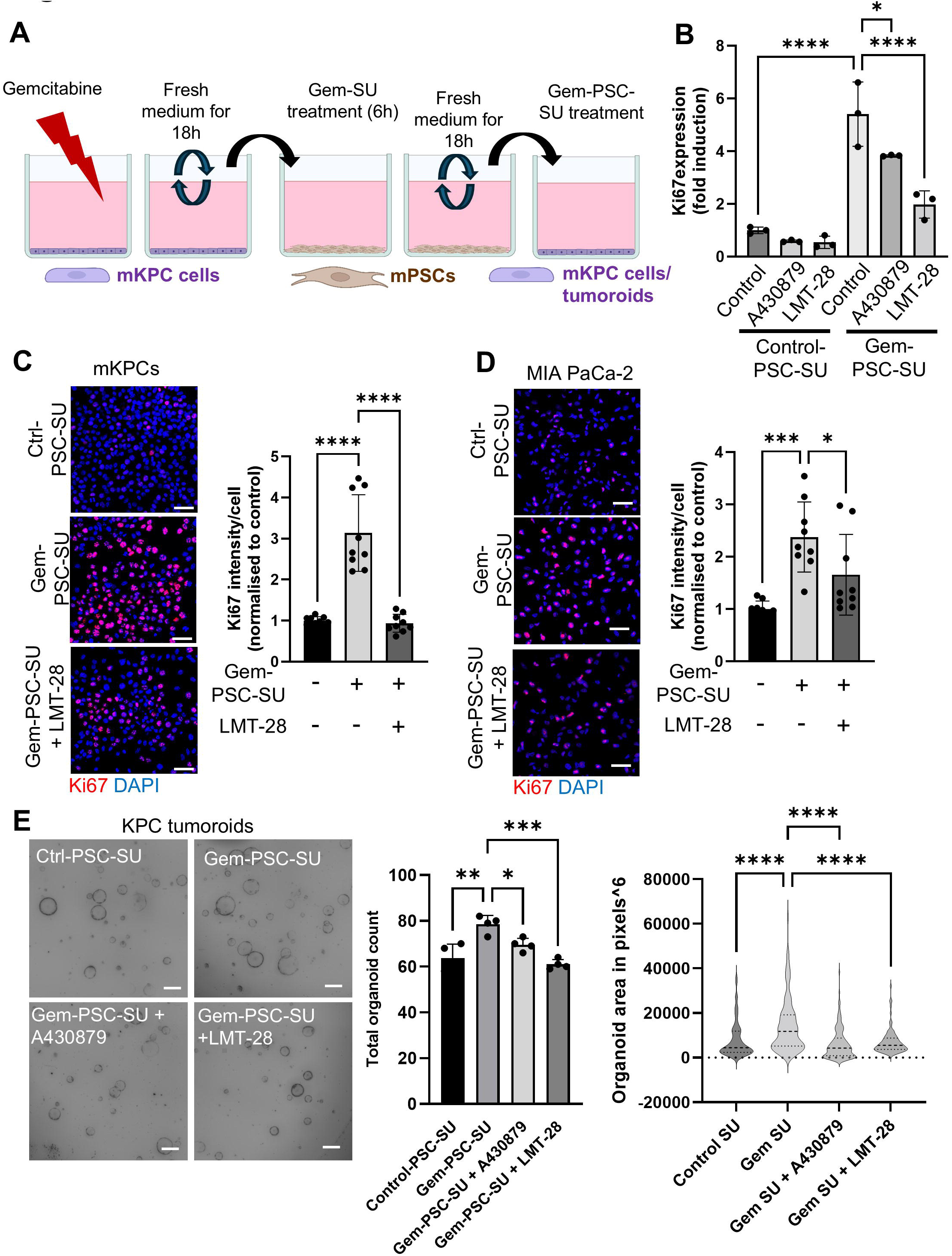
PSCs polarized by dying tumor cells promote cancer cell proliferation via IL-6. **A)** Scheme for the reverse supernatant transfer from Gem-SU treated PSCs on KPC tumor cells or tumoroids. **B)** RT–qPCR analysis of Ki67 in Control-PSC-SU or Gem-PSC-SU treated KPC cells in the absence or presence of the P2X7-inhibitor A430879 or the IL-6R inhibitor LMT-28 (n = 3 biological replicates). **C)** Immunostaining of Ki67 in mKPCs treated with Control-PSC-SU or Gem-PSC-SU in the absence or presence of the IL-6R-inhibitor LMT-28. Nuclei were counterstained with DAPI (scale bar = 100 µm). Graph shows the quantification of Ki67 intensity (pooled data of 3 biological replicates). **D)** Immunostaining of Ki67 in MIA PaCa-2 cells treated with Control-hPSC-SU or Dox-hPSC-SU in the absence or presence of the IL-6R-inhibitor LMT-28. Nuclei were counterstained with DAPI (scale bar = 100 µm). Graph shows the quantification of Ki67 intensity (pooled data of 3 biological replicates). **E)** Representative images of mKPC tumoroids treated as indicated. Graphs show the quantification of the respective tumoroid numbers and sizes (scale bar = 200µm, data were pooled from 2 biological replicates performed in duplicates). All data are mean ± s.d. and were analyzed by one-way ANOVA (B-E) with Tukey’s multiple comparison test (*p≤0.05; **p≤0.01; ***p≤0.0005; **** p≤0.0001; only p values ≤ 0.05 are shown). Illustration in (A) was created using BioRender.com.

### Dying tumor cells educate PSCs to suppress anti-tumor immunity

Inflammatory CAFs are characterized by their high expression of factors that impair anti-tumor immunity (6). In line with this, we detected an upregulation of several genes with known immunosuppressive function in PSCs treated with Gem-SU when compared to PSCs treated with Control-SU (Fig. 4A). To examine whether dying tumor cells modulate the immune regulatory function of PSCs, we co-cultured stimulated T-cells isolated from OT-1 mice (OT-1 cells) together with OVA expressing MC38 cancer cells in Control-PSC-SU or GEM-PSC-SU (Fig. 4B) and measured T-cell-mediated tumor cell killing over time. Strikingly, when co-cultured in Gem-PSC-SU, OT-1 cells were less efficient in killing tumor cells when compared to the co-culture in control medium or Control-PSC-SU (Fig. 4C). When cultured in Gem-PSC-SU, OT-1 cells required not only more time to kill tumor cells upon first contact (Supplementary Data Fig. S4A), they also stopped tumor cell killing after approximately 30 hours of co-culture while T cells cultured with tumor cells in control medium and Control-PSC-SU continued tumor cell killing over the whole 48 hours of analysis (Fig. 4D). Noticeably, around the same time T-cells displayed inhibited killing activity in Gem-PSC-SU, they started to shrink in size, a signal of entering a state of T cell exhaustion (Fig. 4E and F). In line with these observations, flow cytometry analysis confirmed an increase of CD8^+^ T cells expressing the exhaustion markers Tim3, Lag-3^+^ and PD-1 upon exposure to Gem-PSC-SU. Strikingly, this effect was significantly reduced when Gem-PSC-SU were supplemented with the IL-6 receptor antagonist LMT-28, indicating that PSCs polarized by dying tumor cells mediate T cell exhaustion via IL-6 receptor signaling (Fig. 4G and Supplementary Data Fig. S4B).

**Figure 4:**
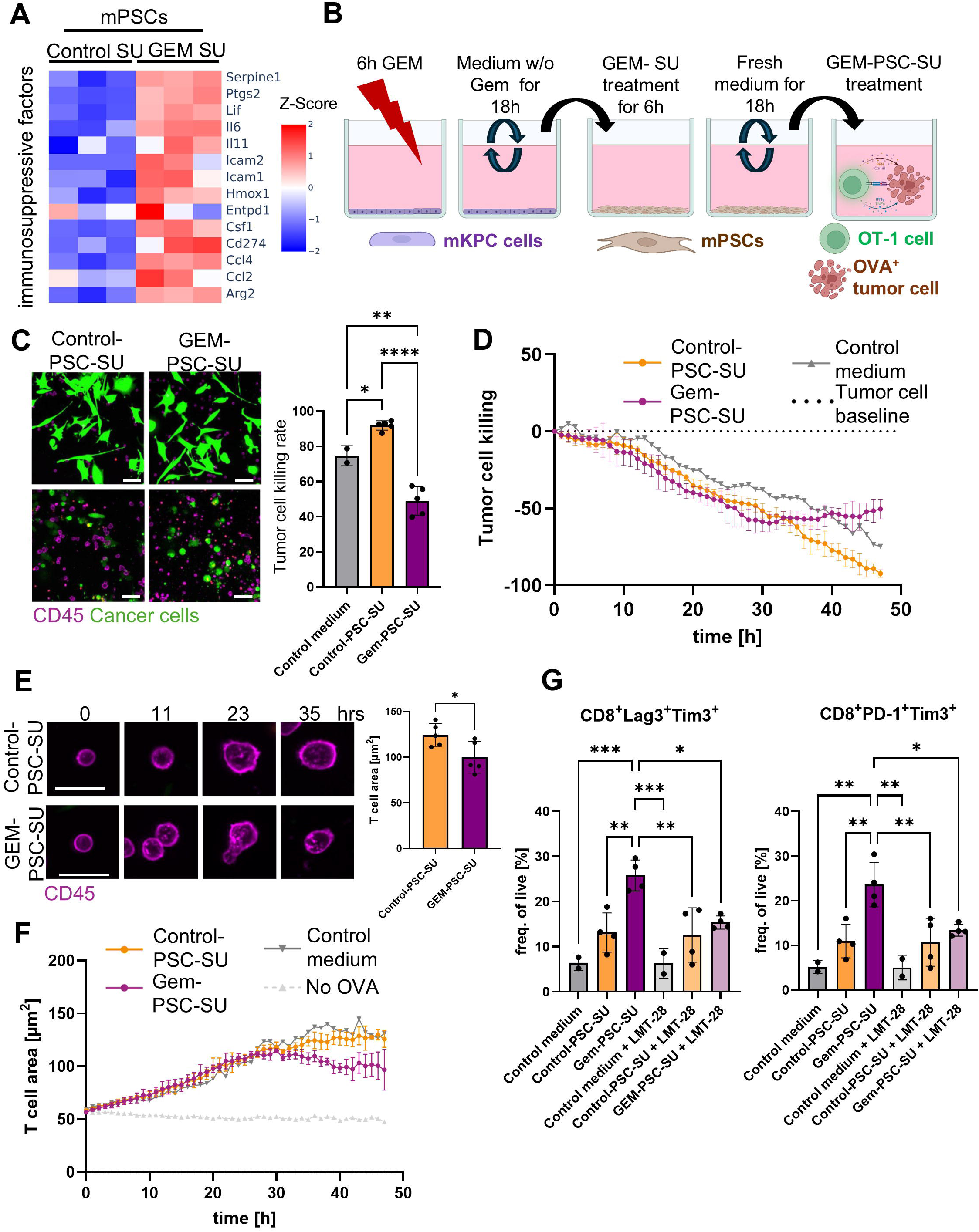
PSCs exposed to factors released from dying tumor cells suppress T cell mediated killing. **A)** RNA-seq analysis of Control-SU- or Gem-SU treated mPSCs. Heatmaps show differentially expressed genes related to immunosuppression. Color codes indicate Z-scores of each condition (n=3 biological replicates). **B)** Scheme for the set-up of the T cell killing assay in the presence of Control-PSC-SU or Gem-PSC-SU. **C)**. Representative images of T-cell/MC38 co-cultures in control T cell medium, Control-PSC-SU or Gem-PSC-SU. Graph shows percentage of killed tumor cells normalized to the number of tumor cells at the start of the experiment (scale bar = 50 µm, n= 3 biological replicates). **D)** Time course analysis of T cell mediated tumor cell killing described in(C). **E)** Representative images and quantification of the area of T cells of the experiment described in (C) (scale bar = 10 µM). **F)** Time course analysis of T cell growth described in(C). **G)** Flow cytometry analysis for the indicated markers of T cells cultured in Control-PSC-SU- or Gem-PSC-SU for 27 hrs. Where indicated the cultured was supplemented with the IL-6 receptor antagonist LMT-28. All data are mean ± s.d. and were analyzed by one-way ANOVA for multiple comparison for the tumor killing and an unpaired t test for the T cell area. (*p≤0.05; **p≤0.01; ***p≤0.0005; **** p≤0.0001; only p values ≤ 0.05 are shown). Illustration in B was created using BioRender.com.

### Pharmacological inhibition of P2X7 improves gemcitabine efficacy against PDAC in vivo

Altogether, our *in vitro* data show that chemotherapy causes the release of factors from dying tumor cells that polarize PSC to tumor-promoting iCAFs, which on the one hand augment tumor cell growth and on the other hand inhibit T cell dependent tumor cell killing. Given these tumor promoting adverse effects of chemotherapeutic intervention, we next addressed whether inhibiting the P2X7-dependent polarization of iCAFs during cytotoxic treatment affects tumor growth *in vivo*. For this we co-injected KPCs together with PSCs subcutaneously in immunocompetent C57BL6/J mice (Fig. 5A) and confirmed presence of both, epithelial cells and fibroblasts in tumor tissues, using antibodies against Epcam and Col1A1, respectively (Fig. 5B). Furthermore, we confirmed expression of P2X7 in tumor tissues, which was more pronounced in stromal cells when compared to PDAC cells (Supplementary Data Fig. S5A). We then treated tumor bearing mice with gemcitabine, the P2X7-inhibitor A438079 or a combination of both. Vehicle treated mice were used as controls. First, we analyzed the tumor tissues 24 hrs post treatment for IL-6 expression via immunostaining. We observed enhanced IL-6 expression in mice that were injected with gemcitabine, showing that chemotherapy promotes IL-6 expression *in vivo* and strikingly, that this effect was reduced in mice treated simultaneously with the P2RX7 antagonist A430879 (Fig. 5 C). Next, we treated mice with subcutaneous tumors for a minimum of two weeks according to the treatment regimen shown in Fig. 5D, in order to examine the effect of the treatments on tumor growth. Here, we observed reduced tumor growth in mice treated with gemcitabine or A430879, the latter in line with previous findings, demonstrating reduced PDAC growth upon P2X7 blockade (20). Compared to the single treatment, the growth rate of tumors was even further reduced in mice treated with a combination of gemcitabine and A430879 (Fig. 5E), demonstrating that P2X7 inhibition increases the efficacy of gemcitabine in our subcutaneous PDAC model. In line with this we also observed less proliferation, fewer viable tumor cells and more necrotic tissues in tumors of mice treated with the combination of gemcitabine and A430879 (Fig. 5F and Supplementary Data Fig. S5B and C), altogether showing that P2X7-inhibition enhances the cytotoxic effect of gemcitabine.

**Figure 5:**
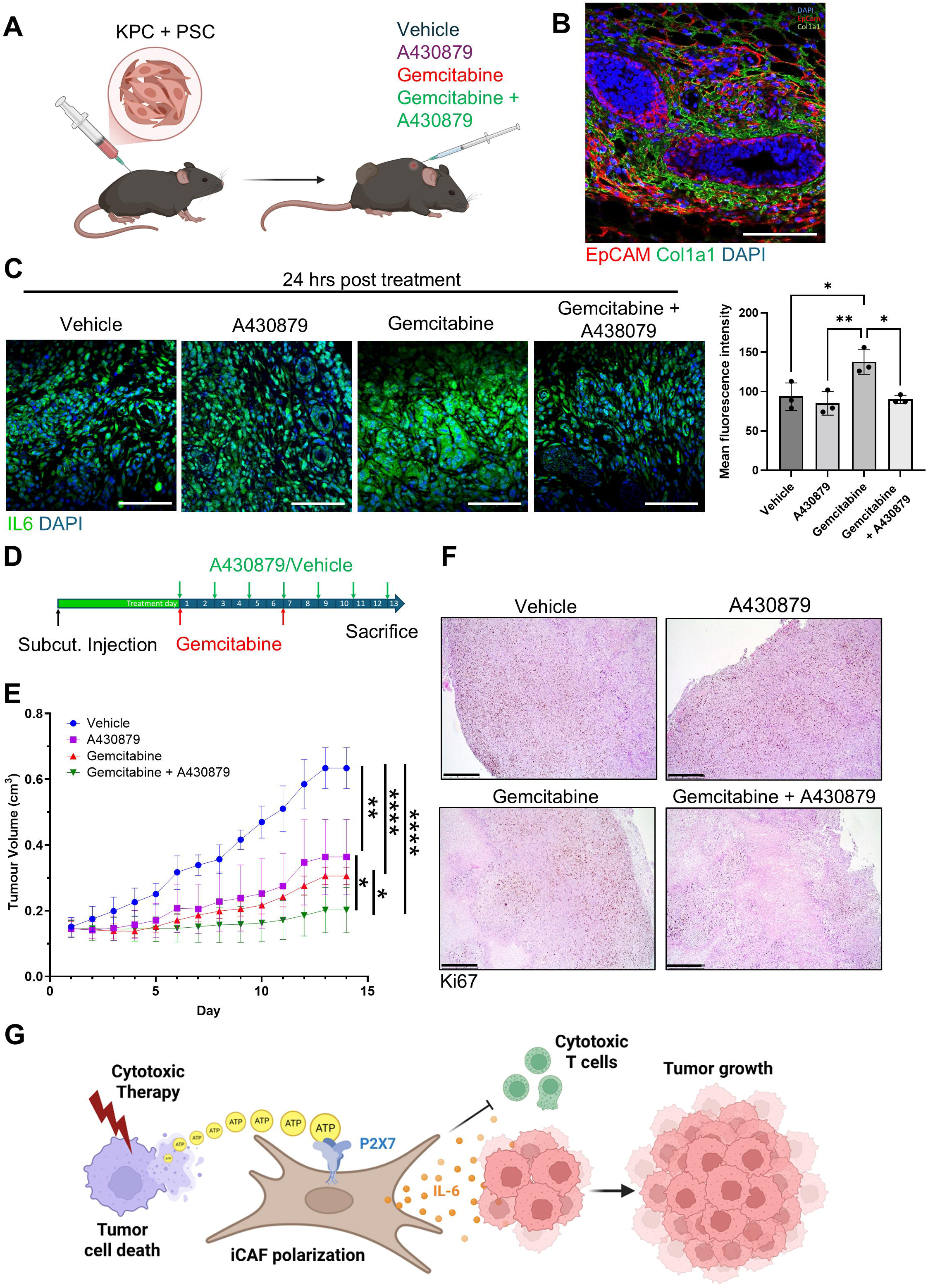
Pharmacological inhibition of P2X7 improves gemcitabine efficacy against PDAC in vivo. **A)** Scheme of subcutaneous PDAC tumor model used in this study. **B)** Immunostaining of subcutaneous tumor tissue section against Epcam and Col1a1 (staining was performed on tumor tissues of 3 independent mice). **C)** IL-6 immunostaining on subcutaneous PDAC tumors 24h post treatment with vehicle, A43087, gemcitabine or a combination of gemcitabine with A430879. Nuclei were counterstained with DAPI (scale bar = 100 µm, n = 5 mice per group). Graph shows quantification of the Il-6 immunofluorescence staining. **D)** Treatment regimen of subcutaneous PDAC tumors analyzed in (E) and (F). **E)** Growth rate of subcutaneous tumors in response to treatment as indicated in (**D**) (n = 7 mice for each group). **F)** Immunostaining of Ki67 in subcutaneous tumors of vehicle treated mice described in (E) (n = 4 independent mice per group, quantification is shown in Supplementary Data Fig. S1G). **G)** Schematic of how dying cells induce IL-6 expression in PSCs to promote tumor cell growth and inhibit anti-tumor immunity. All data are mean ± s.d. and were analyzed by one-way (C) or two-way ANOVA (E) with Tukey’s multiple comparison test (*p≤0.05; **p≤0.01; ****p≤0.0001; only p values ≤ 0.05 are shown).

In summary, we demonstrate that chemotherapy-treated tumor cells release extracellular ATP, which activates P2X7-dependent signaling in PSCs, leading to MEK/ERK activation and robust induction of IL-6 family cytokines. Functionally, these polarized PSCs enhance tumor cell proliferation, induce transcriptional programs associated with epithelial–mesenchymal transition (EMT) and therapy resistance, and impair cytotoxic T cell activity (Fig. 5G). Importantly, pharmacological inhibition of P2X7 significantly improves gemcitabine efficacy *in vivo*, underscoring the therapeutic relevance of this stromal feedback loop.

## Discussion

In this study, we identify a previously unrecognized consequence of cytotoxic therapy in PDAC: the paracrine reprogramming of stromal cells toward a tumor-promoting iCAF phenotype.

The dense desmoplastic stroma of PDAC, long viewed primarily as a physical barrier to drug delivery, is now well-recognized as a highly dynamic and signaling-competent compartment (1). There is a substantial functional heterogeneity and plasticity among CAFs in PDAC, including inflammatory CAFs, myofibroblastic CAFs, and antigen-presenting CAFs (apCAFs)(21). These CAF states are dynamically regulated by microenvironmental cues and can interconvert depending on signaling inputs from tumor and immune cells. Within this framework, our findings identify therapy-induced cancer cell death as potent driver of iCAF polarization, instructing tumor resident fibroblasts to produce protumorigenic and immunosuppressive cytokines, suggesting that chemotherapy—while eliminating therapy-sensitive tumor cells—simultaneously initiates a compensatory stromal program that fosters proliferation and immune evasion of therapy-resistant clones.

Mechanistically, we identify extracellular ATP as a central mediator of this tumor–stroma crosstalk. ATP is a well-established danger-associated molecular pattern (DAMP), released during immunogenic cell death (22). In cancer, purinergic signaling has been implicated in inflammasome activation, immune modulation, tumor cell survival and therapeutic resistance (14, 23). Notably, we have shown previously that cytotoxic therapy triggers the release of ATP from dying tumor cells in order to alert surrounding cancer cells via activation of P2X4 receptors, leading to a rapid increase of tumor cell-intrinsic mTOR-dependent resistance mechanisms (8). Here, we show that this “warning” signal is not restricted to neighbouring epithelial tumor cells but also includes the activation of protumorigenic programs in surrounding stromal cells, however, independent of mTOR.

Instead, we show that ATP derived from dying PDAC cells induces IL-6 production in PSCs in dependence of MEK/ERK signaling, indicating that the downstream response to elevated extracellular ATP levels in the tumor is highly cell-type specific. Although PSCs express multiple purinergic receptor subtypes, by employing selective P2X receptor inhibitors, we identify P2X7 as the potentially dominant receptor driving ERK activation and IL-6 induction in this context and position P2X7 as a critical signaling node linking therapy-induced tumor cell death to stromal reprogramming. Although, we cannot exclude that other P2X receptors are similarly involved, our results are consistent with previous reports showing that ectopic activation of P2X7 receptors in PSCs increases IL-6 release to promote pancreatic cancer cell proliferation (13, 24). We now extend these observations by demonstrating that cytotoxic therapy serves as the upstream trigger for this pathway through cell death–associated ATP release.

Functionally, PSC-derived IL-6 not only enhances tumor cell proliferation and resistance-associated transcriptional programs but also suppresses CD8⁺ T cell–mediated cytotoxicity and induces features of T cell exhaustion. Consequently, therapy-induced iCAF polarization may contribute to both tumor-intrinsic and immune-mediated resistance mechanisms.

Importantly, our *in vivo* experiments reveal that pharmacological P2X7 inhibition attenuates chemotherapy-induced IL-6 expression and significantly enhances gemcitabine efficacy, most likely by mitigating adverse stromal reprogramming. It is important to note that in addition to that, P2X7 inhibition may affect pancreatic tumor growth via other mechanisms, as ATP/P2X7 can also directly promote PDAC cell proliferation and therapeutic resistance (13, 25). Hence, the therapeutic benefit observed with P2X7 blockade likely reflects both, stromal and tumor cell–intrinsic effects.

Given the ongoing efforts to therapeutically target IL-6 signaling in PDAC, the implementation of small-molecule P2X7 inhibitors and recombinant ATP-hydrolyzing ectonucleotidases to interrupt the ATP/P2X7/IL-6 axis may represent a promising therapeutic strategy to improve the efficacy of current treatment regimens (26–28).

Noteworthy, several limitations of our study should be considered. Our *in vivo* experiments were performed in a subcutaneous model, which does not fully recapitulate the anatomical, stromal, and immunological complexity of orthotopic or genetically engineered PDAC models. Furthermore, although ATP emerged as a key mediator, additional DAMPs or soluble factors released from dying tumor cells may cooperate in shaping CAF phenotypes. Although our results highlight a prominent role for IL-6, therapy-educated iCAFs may exert tumor-promoting effects through additional secreted cytokines, including IL-6 family members such as LIF and IL-11 (29), as well as CC chemokine ligands (e.g., CCL2 and CCL4) (30), which we found to be upregulated in PSCs following exposure to supernatants from chemotherapy-treated PDAC cells. Moreover, while we functionally examined the effects of ATP-reprogrammed PSCs on cytotoxic T cells, other cell types within the tumor microenvironment (TME) are likely similarly affected by therapy-induced iCAFs, potentially contributing to the protumorigenic consequences of chemotherapy.

Hence, future studies should explore how purinergic signaling integrates with other inflammatory pathways and define its role within the broader immune landscape of PDAC.

Despite these limitations, our findings support a model in which cytotoxic therapy induces a paradoxical stromal adaptation in PDAC in which ATP released from dying tumor cells reprograms PSCs into IL-6–secreting iCAFs via P2X7 signaling. This adaptive stromal response promotes tumor regrowth and immune suppression, ultimately limiting therapeutic efficacy. Targeting therapy-induced purinergic signaling may therefore represent a promising strategy to enhance chemotherapy responses and overcome microenvironment-driven resistance in PDAC.

More broadly, our findings support a model in which cytotoxic therapy not only eliminates tumor cells but simultaneously generates danger-associated signals that reprogram the tumor microenvironment. In this context, extracellular ATP functions as a therapy-induced paracrine signal that coordinates adaptive tumor–stroma communication, highlighting purinergic signaling as a critical regulator of microenvironment-driven therapeutic resistance in pancreatic cancer.

## Methods

### Animal studies

Mice were kept under SPF conditions in individually ventilated cages with a 12-hour/12-hour light–dark cycle and a standard altromin housing diet. For the subcutaneous tumor models, 8-12-week-old male and female C57BL/6J mice (Charles Rivers Laboratories) were injected with a mixture of 5x10^5 mPSC4 cells and 5x10^5 KPC cells, in a 50% Matrigel (Corning, 354230) and 50% PBS solution into the flanks of the mice while under 2-4% isoflurane (Baxter, HDG9623) narcosis. Tumor sizes were measured with a digital caliper, and tumor volumes were calculated according to the formula Tumour volume (mm^3) = length x width ^2 x 0.52. For studying effects on tumor growth, mice were injected intraperitoneally with saline or 50 mg/kg gemcitabine once per week and where indicated with either 5% DMSO (Carl Roth, A994.2) in saline or 20 mg/kg A 438079 HCL (Selleckchem, S7705) every other day for 2 weeks once tumours reached 150-200 mm^3. Mice were scored and tumours were measured daily, until mice reached the determined endpoints when tumors reached 1.5-1.7 cm^3 or 1.5 cm in diameter. Some mice had to sacrificed previous to this endpoint due to tumor ulceration (mice are indicated in the supplementary data). For studying short term effects of the respective treatments, mice were intraperitoneally injected with 5% DMSO in saline, 20 mg/kg A 438079 HCL, or 50 mg/kg gemcitabine, or a combination of 20 mg/kg A 438079 HCL, or 50 mg/kg 20 mg/kg A 438079 HCL, or 50 mg/kg once tumours reached 1.2-1.5 cm^3 and sacrificed 24 hrs post treatment. At the end of experiments, mice were humanely euthanized to collect tumors. KPC mice have already been described in (31). All studies were approved by the regional agency on animal experimentation (Regierungspräsidium Giessen).

### Cell lines

Mouse pancreatic adenocarcinoma cell lines mKPC4 (in the text referred to as mKPC) were derived from a spontaneous KrasG12D/+, Trp53R172H/+ and Pdx1-Cre (KPC) tumor. The presence of the expected mutations in Kras and Trp53 was verified by Sanger sequencing of PCR-amplified complementary DNAs (cDNA). Mouse pancreatic stellate cell line mPSC4 (referred to as mPSC) and human pancreatic stellate cells hPSC1 (referred to as hPSC) have been published (31–33) and were kindly provided by Albrecht Neesse and Matthias Löhr. MIA PaCa-2 cells obtained from DSMZ (Germany). MC38-OVA murine colon adenocarcinoma cells were kindly provided by Rienk Offringa (DKFZ, Heidelberg, Germany). All cell lines were cultured in Dulbecco’s Modified Eagle Medium (DMEM (high glucose plus glutamine and pyruvate), Invitrogen), supplemented with 10% foetal bovine serum (FBS; Anprotech, Germany), 1% Hepes and 1% penicillin/streptomycin (complete medium) at 37°C with 5% carbon dioxide (CO2). All cells were regularly checked for mycoplasma contamination.

### PDAC tumoroid culture

Murine PDAC organoids were derived from a spontaneous KrasG12D/+, Trp53R172H/+ and Pdx1-Cre (KPC) tumors as previously described (31). Tumoroids were cultured in Matrigel (Corning, 356231) and PDAC organoid medium consisting of advanced DMEM/ F12 (Gibco, 12634028) supplemented with 1% penicillin/ streptomycin (Anprotec, AC-AB--0024), glutamine (Anprotec, AC-AS-0001), 1% HEPES (Anprotec, AC-DS-0007), 50% Wnt3a conditioned medium, 10% R-spondin conditioned medium, 4.7 mg/mL D-glucose (Merck, G7021), 5% Nu-serum IV (Corning, 355104), 0.1 mg/mL Soybean trypsin inhibitor (Thermo Fisher Scientific, 17075029), 25 ug/mL Bovine Pituitary Extract (Corning, 354123), 1% ITS + Premix (Corning, 354352), 100 ng/mL murine EGF (Peprotech, AF-315-09-500µg), 100 ng/mL Cholera Toxin (Sigma-Aldrich, C8052-.5MG), 1 μM Dexamethasone (Sigma-Aldrich, D1756-25MG), 0.5 μM A83-01 (Sigma-Aldrich, SML0788-5MG) and 1.22 mg/mL Nicotinamide (Sigma-Aldrich, N3376). Organoids were maintained in a humidified incubator at 37°C and with 5% CO2, and were passaged twice a week, wherein Matrigel (Corning, 354230) drops were mechanically dissociated via trituration using a glass Pasteur pipette in ice-cold advanced DMEM/F12. The organoid suspension was then centrifuged at 200g for 5 mins reseeding in fresh Matrigel drops which were allowed to solidify at 37°C before adding fresh organoid medium.

### Supernatant transfer experiments

To generate Control-SU and Gem-SU from murine PDAC cells, mKPC4 cells were seeded into 10cm plates at a density of 2.9x10^6 cells/ plate and cultured in complete medium overnight. The next day, medium was replaced with complete medium containing 10 μM gemcitabine (Cayman Chemical, Cay11690) or medium containing equivalent volumes of NaCl. After 6 hrs, medium was removed and cells were washed with PBS (Anprotec, AC-BS-0002). Subsequently, cells were cultured in starvation medium composed of 0.5% FCS, 1% Penicillin/ streptomycin, 1% HEPES. After 184 hrs supernatants were collected from the plates, filtered using a 0,22 µm filter and centrifuged at 700g for 5 mins to remove cellular debris. To generate Control-SU and Dox-SU from human PDAC cells, MIA Paca-2 cells were seeded in 10 cm plates at a density of 4.4x10^6/ plate and cultured in complete medium overnight. The next day, medium was replaced with complete medium containing 5 μM doxorubicin (Hölzel Diagnostika, S1208) for 6 hrs or medium containing equivalent volumes of NaCl. Doxorubicin was used as we observed inefficient cell killing upon gemcitabine treatment. After 6 hrs, medium was removed and cells were washed with PBS (Anprotec, AC-BS-0002). Subsequently, cells were cultured in starvation medium composed of 0.5% FCS, 1% Penicillin/ streptomycin, 1% HEPES. After 18 hrs supernatants were collected from the plates, filtered using a 0,22 µm filter and centrifuged at 700g for 5 mins to remove cellular debris. The supernatants were snap-frozen in liquid nitrogen and stored at - 80°C. In order to generate Control-PSC-SU, Gem-PSC-SU and Dox-PSC-SU from pancreatic stellate cells, murine PSC4 cells and human PSC-1 cells were seeded in a 10cm plate at a density of 8x10^5 cells or 1.4x10^6 cells/plate, respectively and cultured in complete growth medium overnight. The next day, the cells were washed with PBS and incubated for 24h in starvation medium, before culturing the cells with supernatant of vehicle-, gemcitabine- or doxorubicin treated PDAC cells for 6hrs. Where indicated (Fig. 3B and E and Supplementary Data Fig. S3A) P2X7-activation of PSCs was blocked by supplementing tumor cell supernatants with 40 µM A438079 (Santa Cruz Biotechnology, SC-203788B). Following this, the cells were washed with PBS and cultured with fresh starvation medium or organoid medium. After 18 hrs supernatants were collected from the plates, filtered using a 0,22 µm filter and centrifuged at 700g for 5 mins to remove cellular debris. The supernatants were snap-frozen in liquid nitrogen and stored at -80°C. For the treatment of PSCs with Control-, Gem- or Dox-SU, mPSC4 or PSC-1, cells were cultured in complete medium and starved overnight in starvation medium. For RT-qPCR analysis, PSCs were treated for 24 hrs with Control-, Gem- or Dox-SU for immunoblot and immunofluorescence analysis cells were treated for 30 min. Where indicated, supernatants were supplemented with the pharmacological inhibitors rapamycin (10 μM; LC Laboratories, R5000), UO126 (10 μM; LC Laboratories, U6770), PPADS tetrasodium salt (60 μM; Santa Cruz Biotechnology, SC-253332), A438079 hydrochlorides (40 μM, Santa Cruz Biotechnology, SC-203788B) or BAY-1797 (2 μM, Hycultec, HY-130605). For the reverse supernatants transfer from Control-, Gem- and Dox-SU treated PSCs to PDAC cells, mKPC or MIA PaCa-2 cells were cultured in complete growth medium and starved in starvation medium overnight. PDAC cells were then treated with the respective PSC derived supernatant for 24 hours before lysis or fixation for the analysis via RT-qPCR or immunostaining, respectively. Where indicated supernatants were supplemented with 20 μM LMT-28 (MedChemExpress, HY-102084).

### PDAC tumoroid reseeding assay

KPC tumoroids were dissociated as described above and distributed equally in Matrigel for the treatment with Control-PSC-SU or Gem-PSC-SU for 48 hrs. Where indicated PSCs were treated with A430879 (40 µM; Santa Cruz Biotechnology, SC-203788B) while exposed to Gem-SU or the tumoroids were treated with Gem-PSC-SU in the presence of LMT-28 (20 µM; MedChemExpress, HY-102084). Tumoroids were counted and imaged using the Leica Thunder Imager (Leica Microsystems) for the quantification of tumoroid areas.

### RNA Isolation and Real-Time qPCR

RNA was isolated using the QIAwave RNA Mini Kit (Qiagen, 74536) as per the manufacturer’s instructions, including an on-column DNase I (Qiagen, 79254) digestion process for samples used for RNA sequencing analysis. RNA isolated was assessed for purity (A260/A280 ratio >1.6) and concentration by spectrophotometric analysis using the NanoDrop 1000 (PeqLab Biotechnologie GmbH). 500 ng of RNA per sample was used for reverse transcription to cDNA using the RevertAid First Strand cDNA Synthesis Kit (Thermo Fisher Scientific, K1622) with random hexamer primers (Lifetechnologies, SO142), according to manufacturer’s instructions using the GeneAmp PCR System 9700 (Applied Biosystems). RT-qPCR was performed in a volume of 20 μl consisting of 5 ng of diluted cDNA, 1 μM primer (Thermo Fisher Scientific), 50% iTaq Universal SYBR Green Supermix (Bio-Rad, 1725125) and nuclease-free water (Thermo Fisher Scientific, R0582). This reaction mixture was loaded into a 96-well PCR plate in 20 μL/ well in duplicate and run in the qTower3 G RT-qPCR machine (Analytic Jena GmbH). Relative gene expression was determined using Microsoft Excel from the generated Ct values by the 2−ΔΔCt method, with the mean Ct values being normalized to GAPDH. From these values the fold-induction gene expression was determined and plotted. The primers used are listed in Supplementary Table 1.

### Immunoblot analysis

Cells were lysed in Laemmli sample buffer containing 20 mM DTT (Carl Roth, 6908.2), separated by SDS–polyacrylamide gel electrophoresis and transferred to 0.45 μm PVDF membrane (Millipore, number IPVH00010). Membranes were incubated in blocking solution (5% non-fat milk and 1% BSA in PBS) at room temperature for 1 h. After blocking, the membranes were incubated at 4 °C overnight with primary antibodies. After washing with PBS, membranes were incubated with appropriate horseradish peroxidase (HRP)-conjugated secondary antibodies for 1 h at room temperature. After washing in PBS, protein bands were visualized and imaged using enhanced chemiluminescence substrate (ECL) within the ChemiDoc MP Imaging system (Bio-Rad). Immunoblots shown in the manuscript are for representative experiments; the number of successful biological replicates is indicated as *n* in the figure legends. All antibodies used are listed in Supplementary Table 2.

### Immunostaining

For staining of cells, the cells grown on glass coverslips and fixed for 15 mins with 4% paraformaldehyde (PFA) in PBS at room temperature. The fixed cells were washed with PBS and permeabilised with PBST for 10 mins before blocking for 1.5 hrs with a 4% powdered milk (Carl Roth, T145.3) solution in 0.02% Triton X-100 (Carl Roth, 3051.3) in PBS (PBST) at room temperature. Cells were then incubated with primary antibody diluted in blocking buffer at 4°C overnight. The next day, cells were washed with PBST and incubated with secondary antibody and 1 ug/mL DAPI (Sigma Aldrich, D9542-5MG) diluted in blocking buffer for 1 hr at room temperature. The cells were then washed and mounted in aqueous mounting media (Dako, S3023) and stored at 4°C before imaging. For staining of tumor tissues, the tissues were processed for formalin fixation via the HistoCore Pearl tissue processor (Leica Microsystems) After processing tissues were embedded in paraffin using the TES99 Embedding Center (Medite Medical GM) and tissue blocks were cut into 5 μm thick slices using the HistoCore Multicut microtome (Leica Microsystems) which were mounted on positively charged glass slides (Fisher Scientific, 10149870). For staining, these sections were deparaffinised by washing twice in xylene (Carl Roth, 9713.3) for 5 mins followed by rehydration in 100%, 90% and 70% ethanol solutions for 2 mins each before washing in distilled water. Deparaffinized tissue sections underwent antigen retrieval using citrate buffer for 10 mins after boiling and removed after cooling to be washed in ddH2O. Tissues were then permeabilised for 10 mins with 0.2% PBST, blocked for 1.5 hrs with 4% milk in PBST and incubated overnight with a primary antibody diluted in blocking buffer at 4°C. The next morning, the sections were washed with PBST, stained with fluorophore conjugated secondary antibodies and 1 ug/mL DAPI in 4% milk solution before being washed in PBST and mounted with aqueous mounting medium and stored at 4°C. Imaging was performed on the LSM700 confocal microscope with a 20x objective and the images were analysed using ImageJ. Fluorescence intensity was determined by quantified regions of interest. For Ki67 immunohistochemistry, deparaffinized sections were subjected to antigen retrieval, followed by incubation in 3% hydrogen peroxide to quench endogenous peroxidase activity, permeabilization with PBS containing 0.1% Tween-20, and blocking with 5% goat serum. Sections were incubated overnight at 4 °C with anti-Ki67 primary antibody (1:100; BioLegend). After washing in PBS, sections were incubated with a biotinylated secondary antibody (Vector Laboratories) for 1 hour, followed by signal amplification using the VECTASTAIN ABC Kit and visualization with DAB substrate (Vector Laboratories). Slides were mounted using Eukitt Quick-hardening mounting medium (Sigma-Aldrich) and imaged on a Leica DM750 microscope. Ki-67–immunostained sections were digitally analyzed using QuPath (version 0.7.0). Tumor regions were manually annotated, and nuclei were segmented based on the hematoxylin channel. Ki-67 positivity was determined according to nuclear DAB signal intensity. A total of 7–16 images per tumor were analyzed depending on tumor size. Ki-67–positive cell density was calculated as the number of positive cells per mm² of total tumor area.

### RNA sequencing analysis

Murine PSC-4 cells were treated with Control-SU and Gem-SU for 24 hrs, while mKPCs where treated for 24 hrs with Control-PSC-SU or Gem-PSC-SU for 24 hrs and RNA was isolated using the QIAwave RNA Mini Kit (Qiagen, 74536) according to the manufacturer’s instructions, including an on-column DNase I (Qiagen, 79254) digestion process for samples used for RNA sequencing analysis. RNA quality and quantity was evaluated on a 2100 Bio-analyzer (Agilent) using the Agilent RNA 6000 Pico Kit (Agilent Technologies). For RNA sequencing analysis, RNA samples were further processed by Novogene and sequenced via NovaSeq X Plus Sequencing systems (Illumina based). Raw sequencing reads were assessed for quality using FastQC. Adapter sequences and low-quality bases were removed using Trim Galore (v0.6.10), which internally employs Cutadapt (v5.2). Reads were trimmed with a Phred quality score cutoff of 20. Illumina TruSeq adapter sequences were automatically detected and removed. Paired-end reads were processed in paired mode, and read pairs in which at least one mate was shorter than 20 bp after trimming were discarded. The quality of trimmed reads was re-evaluated prior to downstream analysis. RNA-sequencing data were then quantified at the transcript level using Salmon (34). Transcript-level abundance estimates were imported into R (version 4.5.1) and summarized to gene-level counts using tximport (35) based on the Ensembl mouse reference annotation (Mus musculus GRCm39). Differential gene expression analysis was performed using DESeq2 (36), with experimental condition as the main factor. Genes with low expression were filtered prior to analysis. Differential expression was assessed using Wald tests, and p-values were adjusted for multiple testing using the Benjamini–Hochberg method. DESeq2-normalized gene counts were used for visualization of individual gene expression levels and downstream analyses, while variance-stabilized expression values (VST) were used for principal component analysis (PCA) and expression heatmaps. Heatmaps and volcano plots were generated using Flaski (Iqbal et al., 2021 doi: 10.5281/zenodo.4849515).

### Enzyme-Linked Immunosorbent Assay (ELISA)

IL-6 concentration in PSC supernatants was determined using a commercial murine IL-6 ELISA kit (Proteintech, KE10007) as per the manufacturer’s instructions. In brief, recombinant IL-6 protein and supernatants of vehicle and gemcitabine treated PSCs was added to IL-6 antibody pre-coated wells and incubated for 2 hrs at 37°C. The plate was washed with washing buffer, and incubated with 1X detection antibody solution for 1 hr at 37°C. After washing, 1X Streptavidin-HRP solution was added and incubated for 40 mins at 37°C. After washing, TMB substrate solution was added and incubating the plate in the dark for 15 mins before quenching with Stop Solution. Absorbance was measured using the Infinite M Nano plate reader (Tecan Life Sciences) plate reader. IL-6 concentration in supernatant was determined by comparing all values with the values of the provided IL-6 protein standard.

### ATP Measurement and cell viability assay

To determine cell viability and the concentration of ATP in collected dying cell supernatant CellTitre-Glo (Promega, G7570) was used as described in the manufacturer’s instructions. In short, cells were cultured in medium without gemcitabine for 24 hrs. Then cells were treated with medium containing the indicated concentrations of gemcitabine for 6 hrs. After that cells were washed with PBBs and cultured for up to another 18 hrs in medium without gemcitabine. After that cells were lyzed in CellTitre-Glo agent and luminescence was measured as indicated in the protocol. For measuring ATP content in supernatants, the supernatants were collected from KPC4 cells treated with vehicle or 10 μM gemcitabine and mixed 1:1 with the reconstituted CellTitre-Glo reagent in a white opaque 96-well plate. This mixture was agitated using an orbital shaker for 2 mins in the dark before allowing to rest in the dark at room temperature to stabilize the fluorescent signal. Luminescence was then recorded using the Luminoskan Ascent plate reader (Thermo Scientific). Results were plotted against a standard curve generated from measurements of 0.001-1 μM ATP (Sigma-Aldrich, A6419-1G) using Microsoft Excel.

### T cell killing assay

CD3+ T cells were isolated from splenocyte of OT-I transgenic mouse (003831 - OT-1 Strain Details (jax.org)) using the CD8-MACS (Miltenyi Biotec, 130-117-044) according to the manufacturer’s protocol. Target tumor cells (MC38-OVA cell line) were seeded 20 hours before the start of the experiment on fibronectin-coated 96-well imaging plates (Screenstar Microplate, Greiner Bio-One 655866; fibronectin: R&D Systems 1030-FN-05M).

### Live Imaging

On the day of co-culture tumor cells were stained with CellTrackerTM Green CMFDA (Invitrogen, Thermo Fisher, C7025). T cells as well as target cells were suspended in the experimental (60 % conditioned media and 40 % RPMI-1640 (XX) + 10 % FBS (Bio & Sell, FBS.S0615)) and control media (RPMI-1640 (Gibco) + 10 % FBS (Bio & Sell, FBS.S0615)) containing Alexa Fluor® 647 anti-mouse CD45 Antibody (1:500, Biolegend, 103124) for labelling of T cells and NucSpot 568/580 (1:2000, biotium, 41036) for the identification of dead cells. Imaging was conducted with the Cell Voyager CQ1 Confocal Quantitative Image Cytometer (Yokogawa) for 24 hours in intervals of 30 minutes. CellTracker Green was recorded with excitation at 488 nm with 5 % laser power and 500 ms exposure time using a BP525/50 emission filter. NucSpot 568/580 was recorded with excitation at 561 nm at 10 % laser power and 500 ms exposure time using a BP617/73 emission filter. CD45-AF647 was recorded with excitation at 640 nm at 15 % laser power and 500 ms exposure time using a BP685/40 emission filter. Maximum intensity projection images were recorded from three Z planes covering a 3 µm using a 20x/0.8 NA objective (Olympus, UPLXAPO20X). Imaging was performed with four biological replicates of each experimental media in technical duplicates per media condition.

The tumor cell confluency was measured based on the green CellTracker fluorescence using CellPathfinder (version 3.06.01.08, Yokogawa). To calculate the confluency in [%] the total tumor cell area in the field of view area was multiplied * 100.The confluency over time was baseline-corrected and depicted as percentage difference [100*(Value-Baseline)/Baseline]. Baseline-corrected tumor cell confluency = (ConflT[x]-ConflT[0])/ConflT[0] * 100. Each condition was normalized to the growth of the tumor cells without T cell co-culture in the same media. The technical replicates were treated as repeated measures. The CellPathfinder was used to segment T cells and measure their area over time.

### Flow cytometry

CD8+ T effector cells were co-cultured with adherent target cells MC-38 OVA and fixed with Cytofix (BD554722) at different timepoints for flow cytometry staining. After two washing steps with PBS and centrifugation (4 min at 380 g), cells were incubated with titrated fluorochrome-conjugated antibodies in the dark for 20–30 min at 4 °C. After a washing step, the supernatant was discarded and intranuclear staining was performed using the FoxP3/Transcription Factor Staining Buffer Set (Invitrogen 00-5523-00). Cells were resuspended in 100 µl Fix & Perm Cell Permeabilization kit (Invitrogen GAS003) and incubated for 30 min at 4 °C. After two washing steps intracellular antibodies were added and cells were incubated for 30 min at 4 °C. Cells were then washed and measured. Samples were acquired on an LSRFortessa with a 2-Blue, 3-Yellow-Green, 3-Red, 6-Violet, 4-UV configuration, enabling the simultaneous measurement of up to 20 parameters (2 scatter and 18 fluorochromes). Data were saved as FCS files for subsequent analysis using FlowJo v10.10.0.

### Statistical analysis

No statistical method was used to predetermine the sample size for in vivo experiments. For subcutaneous transplantation experiments, mice were distributed among treatment groups when the tumor size reached a measurable size. Tables and graphs for statistical analysis were created using GraphPad Prism 4.03 (GraphPad Software) or Microsoft Excel 2016 MSO (Version 2207 Build 16.0.15427.20166) 32 bit. Statistical significance between two groups was determined by two-tailed Student’s t-test, and that for more than two groups was determined by one-way ANOVA or two-way ANOVA with Tukey’s multiple comparison test (GraphPad Prism 4.03, GraphPad Software). All of the data in the graphs are shown as mean ± s.d., unless stated otherwise.

## Data representation

For the presentation of our data, the figures were created using Microsoft Power Point. Illustrations were created in BioRender (Schmitt, M. (2026) https://BioRender.com/ipx7j7f).

## Data availability

All data relevant to this study are available upon reasonable request. Bulk RNA sequencing described in this study have been deposited in the Gene Expression Omnibus (GEO), and is available using the following identifiers: GSE324779 (bulk RNA sequencing of mKPC and mPSCs). Antibodies and oligonucleotide sequences are listed in Supplementary Table 1 and 2.

## Acknowledgments

This study was financially supported by the Deutsche Forschungsgemeinschaft (DFG; project: KFO325).

## Author contributions

C. M., V. Z., A. O., L. L., A. B., F.A., M.B., designed and performed experiments. M.L. provided KPC tumor tissues and cell lines and reviewed the manuscript. E.S. supervised, designed and analyzed experiments related to immune cells and reviewed the manuscript. C.M. and M.S. wrote the manuscript. M.S. supervised the study and was responsible for concept and design of the study.

---

**Supplementary Data Figure S1 (related to Figure 1).**
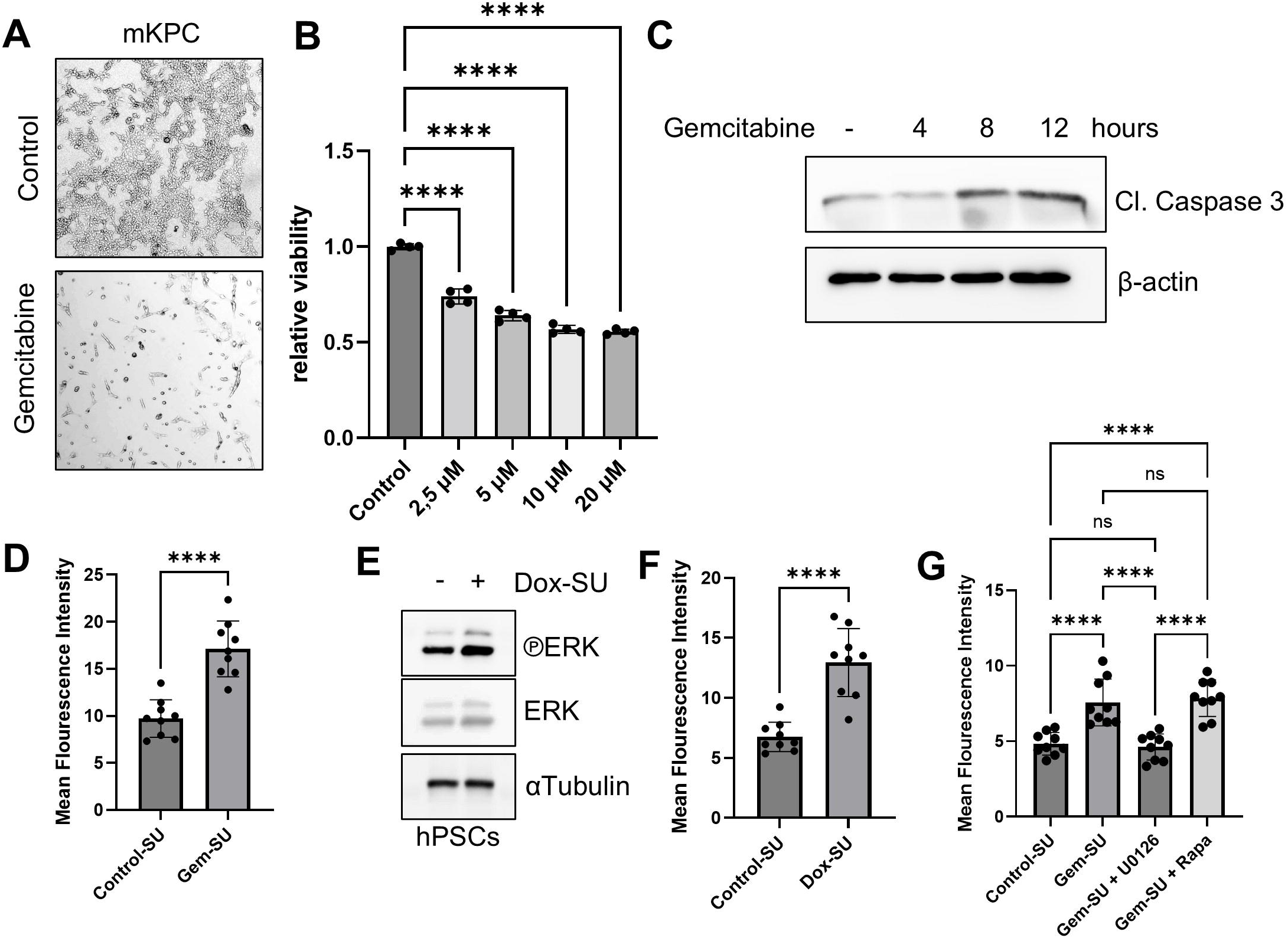
**A)** Bright field images of mKPCs 18h post 6h treatment with Gemcitabine (according to the scheme in Figure 1B). **B)** CellTiter-Glo^®^ cell viability measurement on mPSC treated according to scheme in Fig. 1B, with the indicated concentrations of gemcitabine. **C)** Immunoblots of mPSCs treated with 10 µM gemcitabine for the indicated timepoints (n = 2 biological replicates). **D)** Graph shows the quantification of the mean fluorescence intensity of IL-6 stainings described in Fig. 1H ( pooled data of 3 biological replicates). **E)** Immunoblots of human PSC cells treated for 15 min with supernatants of vehicle (Control-SU) or doxorubicin treated (Dox-SU) MIA PaCa-2 cells (n = 3 biological replicates). **F**) Graph shows the quantification of the mean fluorescence intensity of IL-6 stainings described in Fig. 1J ( pooled data of 3 biological replicates). **G**) Graph shows the quantification of the mean fluorescence intensity of IL-6 stainings described in Fig. 1M ( pooled data of 3 biological replicates). All data are mean ± s.d. and were analyzed by one-way ANOVA with Tukey’s multiple comparison test (****p≤ 0.0001; ns = not significant).

**Supplementary Data Figure S2 (related to Figure 2).**
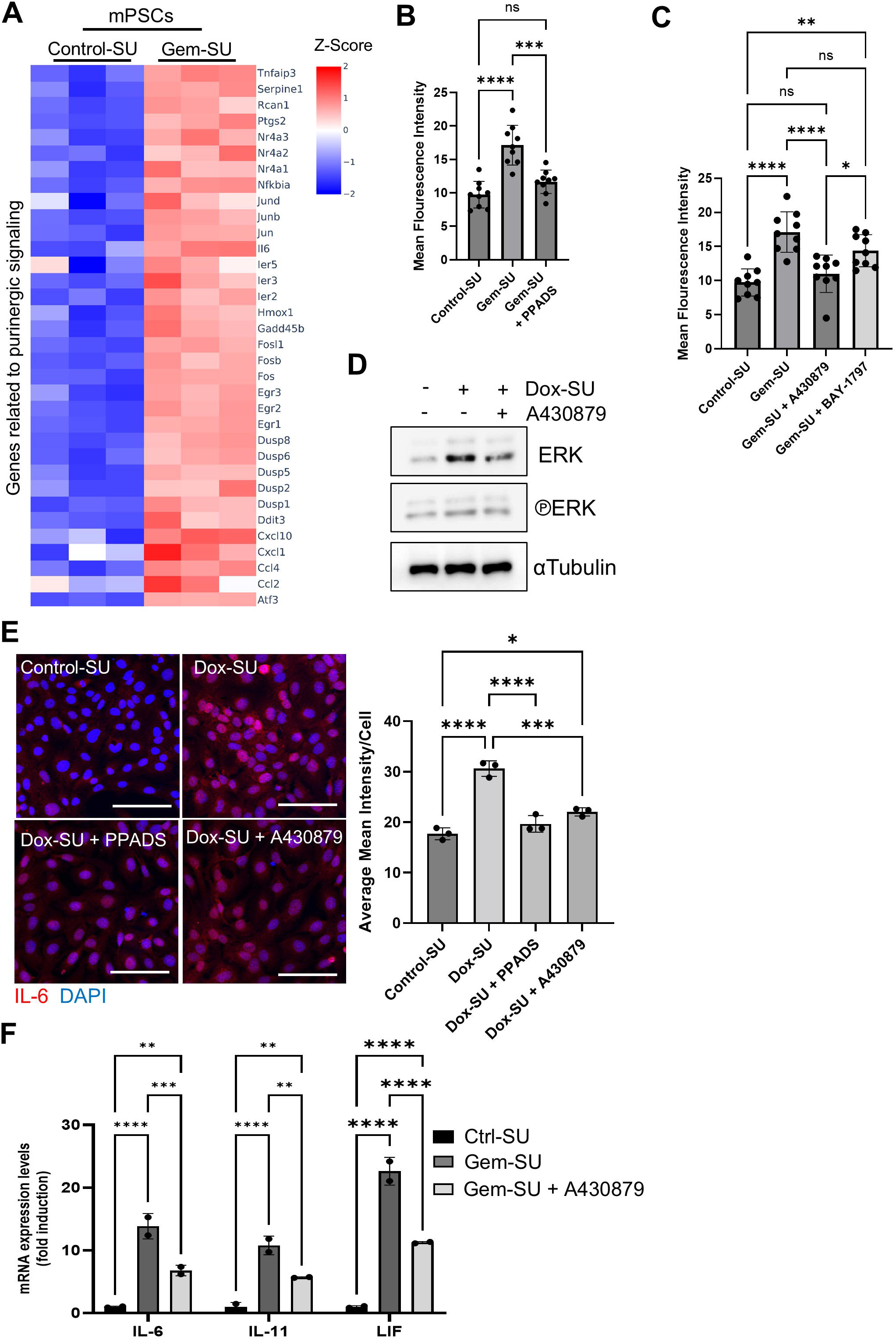
**A)** RNA-seq analysis of Control- or Gem-SU treated mPSCs. Heatmaps show differentially expressed genes related to activation of purinergic signaling. Color codes indicate Z-scores of each condition (n=3 mice per group). **B)** Graph shows the quantification of the mean fluorescence intensity of IL-6 stainings described in Fig. 2G ( pooled data of 3 biological replicates). **C**) Graph shows the quantification of the mean fluorescence intensity of IL-6 stainings described in Fig. 2J (pooled data of 3 biological replicates). **D**) Immunoblots of human PSC cells treated for 15 min with supernatants of vehicle (Control-SU) or doxorubicin treated (Dox-SU) MIA PaCa-2 cells. Where indicated the medium was supplemented with A438079 (n = 3 biological replicates). **E)** IL-6 immunostaining of hPSCs treated with Control- or Dox-SU in the absence or presence of PPADS or A430879 (scale bar = 100 µm, n= 3 biological replicates). Graph shows the quantification of the IL-6-immunofluorescence. **F)** RT–qPCR analysis of the indicated genes in mPSCs treated for 24h with Control-SU or Gem-SU in the absence or presence of the P2X7-inhibtitor A430879 (n = 2 biological replicates). All data are mean ± s.d. and were analyzed by one-way ANOVA with Tukey’s multiple comparison test (*p≤0.05; **p≤0.01; ***p≤0.0005; **** p≤0.0001; ns = not significant).

**Supplementary Data Figure S3 (related to Figure 3).**
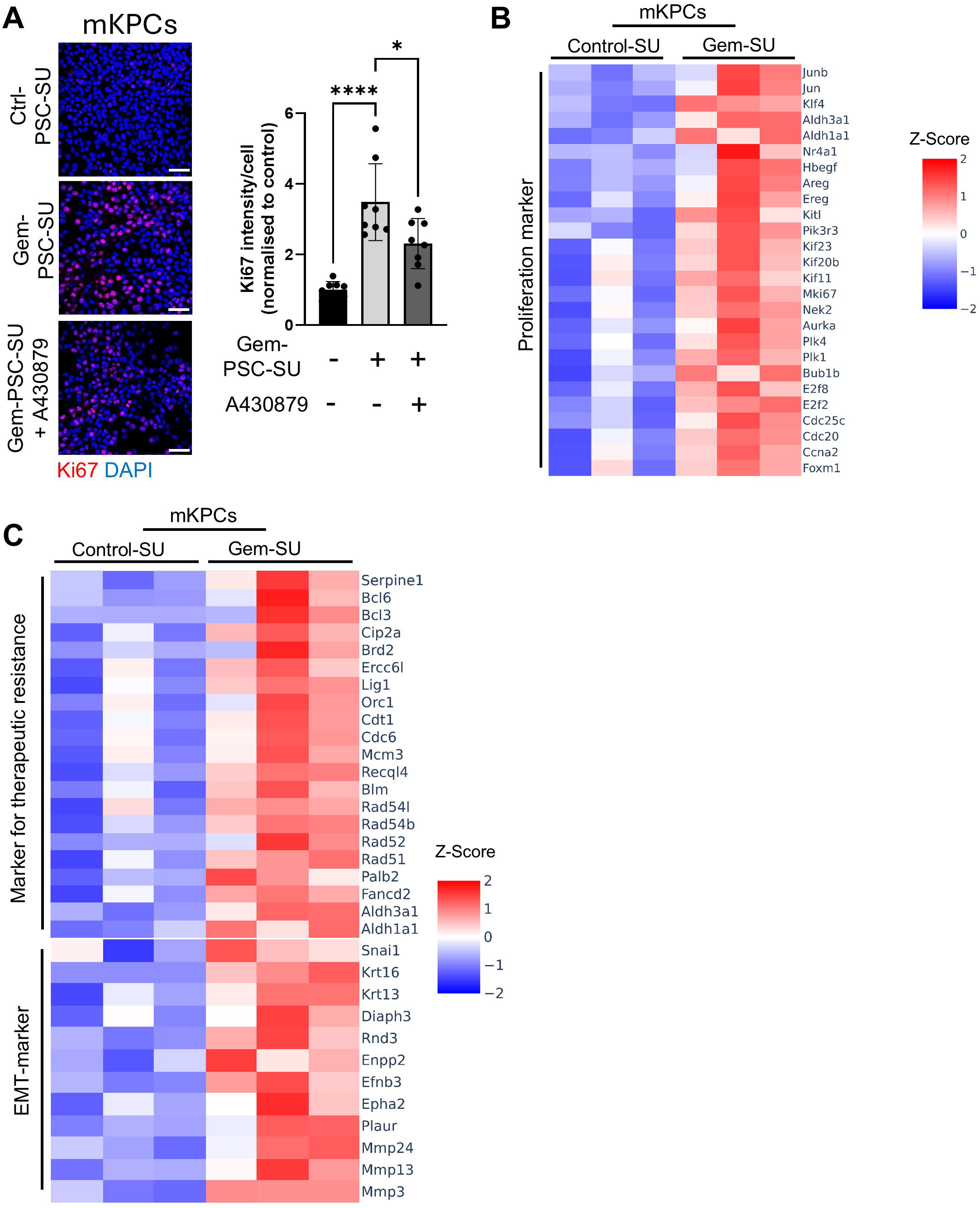
**A)** Immunostaining of Ki67 in mKPCs treated with Control-PSC-SU or Gem-PSC-SU in the absence or presence of the P2X7-inhibitor A430879. Nuclei were counterstained with DAPI (scale bar = 100µm). Graph shows the quantification of Ki67 intensity (pooled data of 3 biological replicates). **B** and **C)** RNA-seq analysis of Control-PSC-SU- or Gem-PSC-SU treated mKPCs. Heatmaps show differentially expressed genes related to proliferation (B), therapeutic resistance and EMT (C). Color codes indicate Z-scores of each condition (n=3 mice per group). All data are mean ± s.d. and were analyzed by one-way ANOVA with Tukey’s multiple comparison test (*p≤0.05; **** p≤0.0001; only p values ≤ 0.05 are shown).

**Supplementary Data Figure S4 (related to Figure 4).**
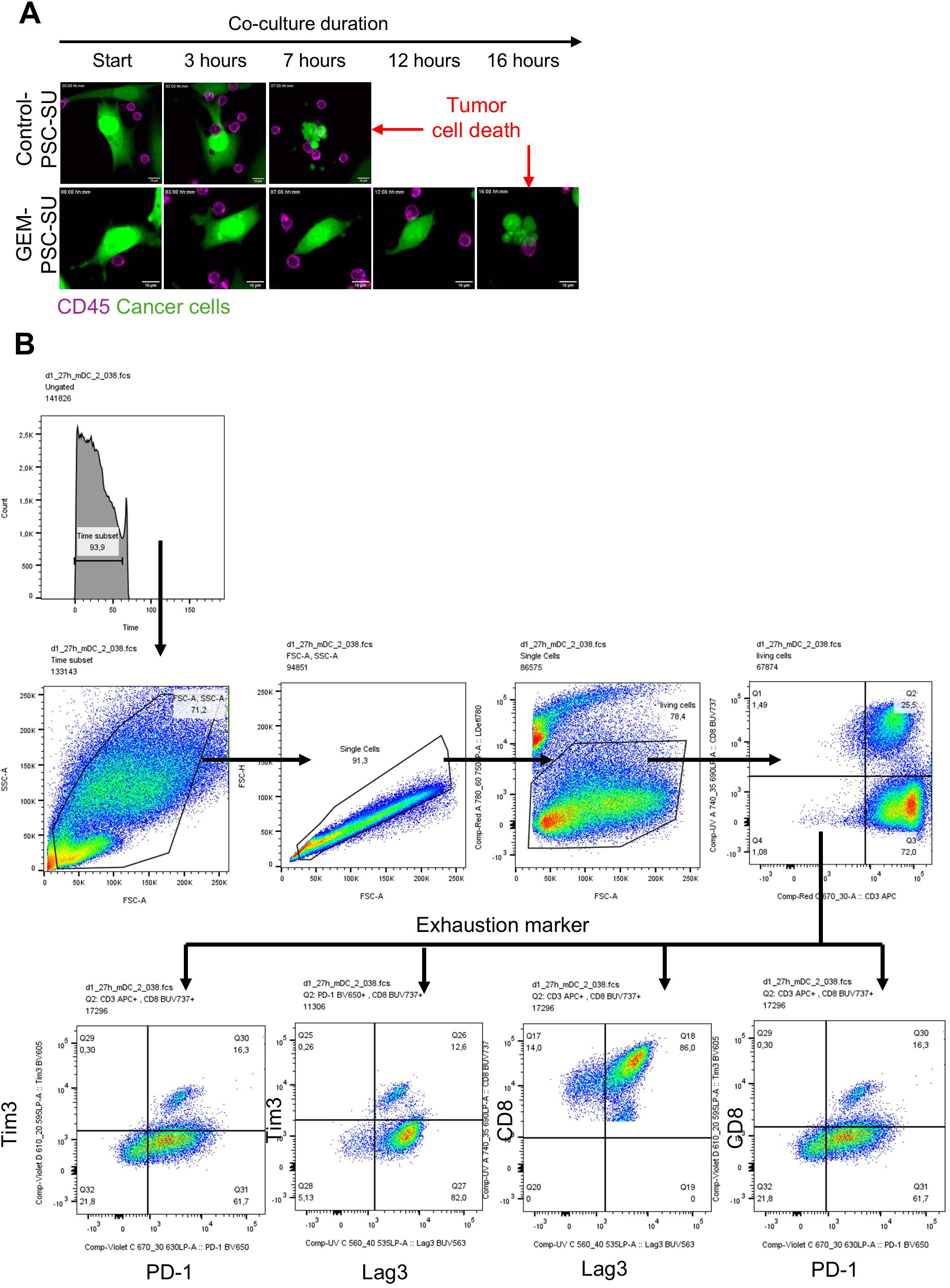
**A)** Analysis of the first killing experiment in T-cell/MC38 co-cultures in Control-PSC-SU or Gem-PSC-SU (analysis was performed on experiments described in Fig. 4C. **B)** Gating strategy of the FACS analysis described in Fig. 4F.

**Supplementary Data Figure S5 (related to Figure 5).**
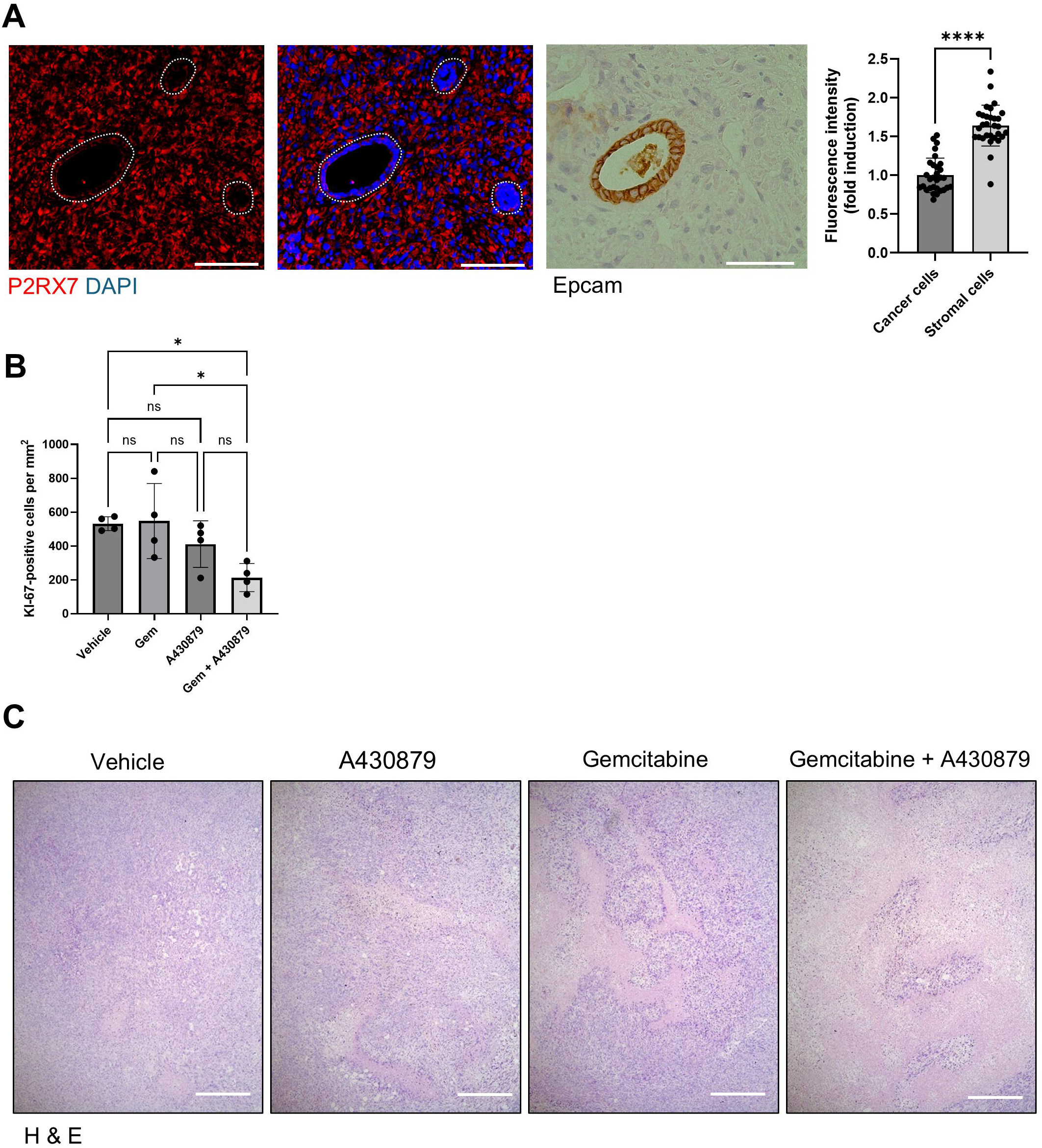
**A)** Immunostaining of P2X7 and Epcam in subcutaneous tumors of vehicle treated mice described in Fig. 5 C). Nuclei were counterstained with DAPI (scale bar = 100µm, n = 3 biological replicates). Graph shows the quantification of the mean fluorescence intensity of P2X7 staining in stromal and PDAC cells. **B)** Graph shows quantification of Ki67 positive cells in tumor tissues described in Fig. 5F (n = 4 independent mice per group). **C)** H&E analysis of tumors analyzed in Fig. 5E (n = 4 independent mice per group). All data are mean ± s.d. and were analyzed by two-tailed Student’s *t*-test ((*p≤0.05; ****p≤0.0001; ns = not significant).

