## Supplementary material for "Tumor Cell Death Drives Tumor-Promoting IL-6⁺ iCAF formation via P2X7-activation": Suplementary Table 1

### List of antibodies used in this study

### A) Antibodies used for immunoblot and immunostaining

| Specificity | Company | Cat. Number | Dilution – cell IF | Dilution – tissue IF/ IHC | Dilution – Immunoblot |
|---|---|---|---|---|---|
| β-Actin | Cell Signalling Technologies | 4970S | - | - | 1:1000 |
| CD8 alpha | Cell Signalling Technologies | 98941S | - | 1:200 | - |
| CD274 (PD-L1, B7-H1) | Biolegend | 124302 | 1:100 | - | - |
| Col1a1 | Cell Signalling Technologies | 72026S | - | 1:200 | - |
| CD326 (EpCAM) | Thermo Fisher Scientific | 14-5791-85 | - | 1:100 | - |
| Cleaved Caspase 3 | Cell Signalling Technologies | 9661S | - | - | 1:500 |
| GAPDH | Cell Signalling Technologies | 2118S | - | - | 1:1000 |
| IL-6 | AB Clonal | A0286 | 1:200 | 1:100 | - |
| Ki-67 | BioLegend | 151202 | 1:200 | 1:100 | - |
| P2X7 Receptor (extracellular) | Almone Labs | APR-008 | 1:200 | 1:100 | - |
| p42/44 | Cell Signalling Technologies | 4695S | - | - | 1:1000 |
| phospho p44/42 (Erk 1/2) | Cell Signalling Technologies | 4370S | - | - | 1:2000 |
| phospho-S6 Ribosomal Protein | Cell Signalling Technologies | 4858S | - | - | 1:2000 |
| S6 Ribosomal Protein | Cell Signalling Technologies | 2217S | - | - | 1:1000 |
| α-Tubulin | Cell Signalling Technologies | 2125S | - | - | 1:2000 |

| | | | | | |
|---|---|---|---|---|---|
| Alexa Fluor® 647 anti-mouse CD45 Antibody | Biolegend | 103124 | 1:500 | - | - |
| Donkey anti-Rabbit IgG (H+L) Highly Cross-Adsorbed Secondary Antibody, Alexa Fluor™ 594 | Invitrogen | A-21207 | 1:400 | 1:400 | - |
| Goat anti-Rat IgG (H+L) Cross-Adsorbed Secondary Antibody, Alexa Fluor™ 555 | Themo Fisher Scientific | A-21434 | 1:400 | 1:400 | - |
| Goat anti-Rabbit IgG (H+L) Cross-Adsorbed Secondary Antibody, Alexa Fluor™ 488 | Scientific | A-11008 | 1:400 | 1:400 | - |
| Goat Anti-Rabbit IgG (H + L)-HRP Conjugate | Bio-Rad | 1706515 | - | - | 1:4000 |

**B) Antibodies and fluorochromes used for flow cytometry**

| Specificity | Company | Catalogue Number | Dilution |
|---|---|---|---|
| Fixable Viability Dye eFluor™ 780 | eBioscience | 65-0865-14 | 1:1000 |
| BD Horizon™ BUV737 Rat Anti-Mouse CD8a | BD Biosciences | 612759 | 1:100 |
| Brilliant Violet 605™ anti-mouse CD366 (Tim-3) Antibody | Biolegend | 119721 | 1:100 |
| BUV563 Rat Anti-Mouse CD223 (LAG-3) | BD Biosciences | 741350 | 1:100 |
| BD OptiBuild™ BV650 Rat Anti-Mouse CD279 (PD-1) | BD Biosciences | 748266 | 1:25 |
