## Supplementary Table 2 for "Tumor Cell Death Drives Tumor-Promoting IL-6⁺ iCAF formation via P2X7-activation"

List of primers used in this study:

| Gene Target | Sequence |
|---|---|
| Murine Il-6-f | CTGGTGACAACCACGGCCTTCCCTA |
| Murine Il-6-r | ATGCTTAGGCATAACGCACTAGGTT |
| Murine Cd274-f | TGCGGACTACAAGCGAATCACG |
| Murine Cd274-r | CTCAGCTTCTGGATAACCCTCG |
| Murine Cd28-f | CACCTCTGATGGAAGCAGCTTC |
| Murine Cd28-r | GATAATCGCTGGTCAGAGCTTCG |
| Murine Cd80-f | CCTCAAGTTTCCATGTCCAAGGC |
| Murine Cd80-r | GAGGAGAGTTGTAACGGCAAGG |
| Murine Cd86-f | ACGTATTGGAAGGAGATTACAGCT |
| Murine Cd86-r | TCTGTCAGCGTTACTATCCCGC |
| Murine Ctla-4-f | GTACCTCTGCAAGGTGGAACTC |
| Murine Ctla-4-r | CCAAAGGAGGAAGTCAGAATCCG |
| Murine Pd1-f | CGGTTTCAAGGCATGGTCATTGG |
| Murine Pd1-r | TCAGAGTGTCGTCCTTGCTTCC |
| Murine Il-11-f | CTGACGGAGATCACAGTCTGGA |
| Murine Il-11-r | GGACATCAAGTCTACTCGAAGCC |
| Murine Il-33-f | CTACTGCATGAGACTCCGTTCTG |
| Murine Il-33-r | AGAATCCCGTGGATAGGCAGAG |
| Murine Cxcl1-f | TCCAGAGCTTGAAGGTGTTGCC |
| Murine Cxcl1-r | AACCAAGGGAGCTTCAGGGTCA |
| Murine Cxcl12-f | GGAGGATAGATGTGCTCTGGAAC |
| Murine Cxcl12-r | AGTGAGGATGGAGACCGTGGTG |
| Murine Lif-f | TCAACTGGCACAGCTCAATGGC |
| Murine Lif-r | GGAAGTCTGTCATGTTAGGCGC |
| Murine Tnfa-f | CAGGAGGGAGAACAGAAACTCCA |
| Murine Tnfa-r | CCTGGTTGGCTGCTTGCTT |
| Murine Il-1a-f | ACGGCTGAGTTTCAGTGAGACC |
| Murine Il-1a-r | CACTCTGGTAGGTGTAAGGTGC |
| Murine Il-1b-f | GGGTCCGTCAACTTCAAAGA |
| Murine Il-1b-r | TGAAGCAGCTATGGCAACTG |
| Murine Il-10-f | CGGGAAGACAATAACTGCACCC |
| Murine Il-10-r | CGGTTAGCAGTATGTTGTCCAGC |

| | |
|---|---|
| Murine Ifna-f | TTTCCCCTGACCCAGGAAGATG |
| Murine Ifna-r | CTCTCAGTCTTCCCAGCACATT |
| Murine Ilfy-f | CAGCAACAGCAAGGCGAAAAAGG |
| Murine Ilfy-r | TTTCCGCTTCCTGAGGCTGGAT |
| Murine Tgfb1-f | TGATACGCCTGAGTGGCTGTCT |
| Murine Tgfb1-r | CACAAGAGCAGTGAGCGCTGAA |
| Murine Hgf-f | GTCCTGAAGGCTCAGACTTGGT |
| Murine Hgf-r | CCAGCCGTAAATACTGCAAGTGG |
| Murine Mki-67-f | GAGGAGAAACGCCAACCAAGAG |
| Murine Mki-67-r | TTTGTCCTCGGTGGCGTTATCC |
| Murine Myc-f | TCGCTGCTGTCCTCCGAGTCC |
| Murine Myc-r | GGTTTGCCTCTTCTCCACAGAC |
| Murine aSma-f | TTCGTGTGGCCCCTGAAGAGCAT |
| Murine aSma-r | CCAGTTGTACGTCCAGAGGCA |
| Murine Vimentin-f | CGGAAAGTGGAATCCTTGCAGG |
| Murine Vimentin-r | AGCAGTGAGGTCAGGCTTGGAA |
| Murine Demin-f | GCGGCTAAGAACATCTCTGAGG |
| Murine Demin-r | ATCTCGCAGGTGTAGGACTGGA |
| Murine Fn1-f | AGTGACAGCATACAGGGTGATGG |
| Murine Fn1-r | CTACGGAGAGACAGGAGGAAATAGC |
| Murine Fap-f | CCGCGTAACACAGGATTCACTG |
| Murine Fap-r | CACACTTCTTGCTCGGAGGAGA |
| Murine G-fap-f | CACCTACAGGAAATTGCTGGAGG |
| Murine G-fap-r | CCACGATGTTCCTCTTGAGGTG |
| Murine Col1a-f | CCTCAGGGTATTGCTGGACAAC |
| Murine Col1a-r | CAGAAGGACCTTGTTTGCCAGG |
| Murine P2x1-f | CATGGGGACAGCTCCTTTGT |
| Murine P2x1-r | GAGTGCAGCCACTGTCATCT |
| Murine P2x2-f | CCAAGGCACCCCTCAAGTAG |
| Murine P2x2-r | CTCTGCCCCTTCTCCCAAAG |
| Murine P2x3-f | TGCTTCAACCAACCCAGTGT |
| Murine P2x3-r | TAAGAGCCCCTCTTCTCCCC |
| Murine P2x4-f | CCTGGCTTACGTCATTGGGT |
| Murine P2x4-r | AAGTGTTGGTCACAGCCACA |

| | |
|---|---|
| Murine P2x5-f | TCTACTGCCCCATCTTCCGA |
| Murine P2x5-r | ATAGTGTGGGTTGCAGTGGG |
| Murine P2x6-f | GCTGCACCATGGACCTACTT |
| Murine P2x6-r | GCTTCAGGTGAGCTGTTCCT |
| Murine P2x7-f | GCACGAATTATGGCACCGTC |
| Murine P2x7-r | CCCCACCCTCTGTGACATTC |
| Murine P2y1-f | TTATGTCAGCGTGCTGGTGT |
| Murine P2y1-r | ACGTGGTGTCATAGCAGGTG |
| Murine P2y2-f | TCAAACCGGCTTATGGGACC |
| Murine P2y2-r | GGCAGCTGAGGTCAAGTGAT |
| Murine P2y4-f | GCTCTATCTGTTCACGGGGG |
| Murine P2x4-r | AGGGAGGAAGCAGTTGTTCG |
| Murine P2x6-f | GGGTAGTGTGTGGAGTCGTG |
| Murine P2x6-r | AGCGAGTAGACAGGATGGGT |
| Murine P2x12-f | TGCTGTACACCGTCCTGTTC |
| Murine P2x12-r | CGGCTCCCAGTTTAGCATCA |
| Murine P2x13-f | GCATCAGGTGGTCAGTCACA |
| Murine P2x13-r | GTGGGGCAAAGCAGACAAAG |
| Murine P2x14-f | CCACATTGCCAGAATCCCCT |
| Murine Gapdh-f | AGACGGCCGCATCTTCTTGT |
| Murine Gapdh-r | TGCCGTTGAATTTGCCGTGA |
| Murine P2x14-r | AGCCGAGAGTAGCAGAGTGA |
| Human Il-6-f | AGACAGCCACTCACCTCTTCAG |
| Human Il-6-r | TTCTGCCAGTGCCTCTTTGCTG |
| Human GAPDH-f | AGCCACATCGCTCAGACAC |
| Human GAPDH-r | GCCCAATACGACCAAATCC |
